# Smart Dura: a functional artificial dura for multimodal neural recording and modulation

**DOI:** 10.1101/2025.02.26.640369

**Authors:** Sergio Montalvo Vargo, Nari Hong, Tiphaine Belloir, Noah Stanis, Jasmine Zhou, Karam Khateeb, Gaku Hatanaka, Zabir Ahmed, Ibrahim Kimukin, Devon J. Griggs, Wyeth Bair, Azadeh Yazdan-Shahmorad, Maysamreza Chamanzar

## Abstract

A multi-modal neural interface capable of long-term recording and stimulation is essential for advancing brain monitoring and developing targeted therapeutics. Among the traditional electrophysiological methods, micro-electrocorticography (μECoG) is appealing for chronic applications because it provides a good compromise between invasiveness and high-resolution neural recording. When combining μECoG with optical technologies, such as calcium imaging and optogenetics, this multi-modal approach enables the simultaneous collection of neural activity from individual neurons and the ability to perform cell-specific manipulation. While previous efforts have focused on multi-modal interfaces for small animal models, scaling these technologies to larger brains, of primates, remains challenging. In this paper, we present a multi-modal neural interface, named Smart Dura, a functional version of the commonly used artificial dura with integrated electrophysiological electrodes for large cortical area coverage for the NHP brain. The Smart Dura is fabricated using a thin-film microfabrication process to monolithically integrate a micron-scale electrode array into a soft, flexible, and transparent substrate with high-density electrodes (up to 256 electrodes) while providing matched mechanical compliance with the native tissue and achieving high optical transparency (exceeding 97%). Our *in vivo* experiments demonstrate electrophysiological recording capabilities combined with neuromodulation, as well as optical transparency via multiphoton imaging. This work paves the way toward a chronic neural interface that can provide large-scale, bidirectional interfacing for multimodal and closed-loop neuromodulation capabilities to study cortical brain activity in non-human primates, with the potential for translation to humans.

## Introduction

Large-scale multi-modal neural recording and manipulation across different areas of the brain with high spatiotemporal resolution, especially on larger brains of non-human primates (NHPs), is crucial to understanding the neural basis of function and dysfunction and translating those findings to humans. While there has been significant recent effort to design such systems for multi-modal high-density recording and stimulation in small animal models, particularly mice^1–5^, there have only been limited interfaces for large-scale neural recording and manipulation in NHPs. Both from the technology and the application standpoint, the neural interfaces designed for rodents cannot be readily translated into NHPs. Developing chronic multi-modal high-resolution neural interfaces for NHPs that can cover multiple areas of the brain with minimal damage to the neural tissue has been an outstanding need.

Leveraging advanced microfabrication techniques, micro-electrocorticography (μECoG) arrays have been developed to record local field potentials, single- and multi-units from the brain surface while remaining minimally invasive^6–8^. Biocompatible, flexible polymers such as polyimide or parylene C have been used to implement μECoGs with improved mechanical compatibility with neural tissue. Still, these devices exhibit a mechanical mismatch with the native dura, thus compromising their potential for chronic stability and rendering them mostly limited to short-term acute experiments^9^.

On the other hand, artificial dura, a transparent and passive ‘cap’ made of soft polydimethylsiloxane (PDMS) elastomers, has proven to be a chronically viable replacement for the native dura to cover open craniotomies, allowing optical access for optical stimulation and recording^10,11^. A few groups have tried to combine artificial dura with ECoG arrays, but either the artificial dura was temporarily removed to allow for the placement of the arrays for the duration of neurophysiology experiments^12^ or it was heterogeneously combined with the ECoG array for chronic placement^13^. The former allowed electrophysiology from anesthetized and awake NHPs with simultaneous optical access, however, the placements and replacements over time can potentially cause increased tissue growth and damage, compromising the stability of the interface. In the latter, heterogeneous methods have accelerated tissue growth between the artificial dura and ECoG array, obstructing any light penetration to the brain.

Homogeneous integration of electrodes on soft elastomer has been demonstrated in spinal cord implants with desired flexibility and stability^14^, showing the capabilities of elastomer-based electrode arrays. More recently, Griggs et al. demonstrated the use of a commercially available ECoG array molded in an artificial dura^15^, named multi-modal artificial dura (MMAD), for simultaneous electrophysiology and optical access where they reported stable optical access (with simultaneous neural recording) similar to the passive artificial dura. However, their nanoparticle ink printing fabrication technology could only provide a limited resolution of 32 electrodes across 3 cm^2^ of the cortex and large-sized opaque electrodes (0.5 mm in diameter) and traces (250-300 μm in width) that limited optical access^15,16^.

The promises of homogenous integration dovetail with a pressing need for high-density monolithically integrated ECoGs and artificial dura using micro-fabrication methods on softer and transparent substrates^17^, such as PDMS. PDMS is a soft, transparent, and biocompatible elastomer, but is not easily implemented in fabrication processes for large-scale production of soft electronics. For example, thin film microfabrication technology at the wafer scale relies on high-resolution photolithography techniques for parallel, high-throughput fabrication. However, due to its low surface energy, PDMS cannot be wetted with photoresist, and therefore, it is not directly compatible with photolithography. Additionally, microfabrication processes rely on rigid wafers for mechanical support during the carry and transfer of the samples to different fabrication tools. Thus, PDMS needs to be applied onto a silicon or glass carrier wafer for microfabrication, but the strong adhesion between PDMS and the carrier wafer makes the release for a standalone PDMS substrate impossible if applied on its own. Additional fabrication steps, such as those implemented in Ji et al.^18^ and others^19,20^, are needed to circumvent the issues with PDMS and make this material compatible with microfabrication processes for biomedical implants.

Achieving micron-size features for electrodes and traces on soft and transparent substrates will not only improve the mechanical properties for chronic stability but will also significantly increase the level of transparency as there is much less opaque material blocking light. Higher transparency can improve the compatibility with optical methods for interrogating the brain. Optical methods such as calcium imaging and optogenetics, which provide cell-type specific recording with single-cell resolution and manipulation of neurons with millisecond temporal precision^21^, are used in conjunction with electrophysiological techniques to study the brain across different spatiotemporal scales for a more comprehensive understanding^22^. However, hybrid technologies that provide electrophysiological and optical capabilities in one platform have been demonstrated mainly in small compact form factors better suited for small animal models^22,23^. There is still an outstanding need to scale up multi-modal neural interfaces to study larger brains, such as those of NHPs, as they provide greater translational potential to humans.

In this paper, we discuss the design and demonstration of microfabricated μECoG array directly implemented on a PDMS artificial dura to provide very high-resolution surface electrophysiology capability as well as optical access. We demonstrate use-case scenarios for multi-modal neural interfacing in macaque brains through local neural recording in several cortical regions during behavior or with simultaneous tactile, electrical, and optogenetic stimulation as well as optical imaging using two-photon microscopy.

## Results

### Smart Dura: concept, design and fabrication

Here we introduce Smart Dura, a high-density surface electrode array fabricated on transparent, soft, and flexible polymer substrates that can seamlessly replace the native dura while enabling optical access to the brain (Fig. 1). This neural interface, implemented on a stack of Parylene C and PDMS, two biocompatible flexible polymers, is carefully designed to match the mechanical properties of native dura. Therefore, it can replace the native dura and be used as a chronic long-term replacement, with the unique advantages that (i) it provides high-resolution electrophysiological recording and electrical stimulation; and (ii) it is transparent and thus provides optical access to the neural tissue for multi-modal optical imaging and neuromodulation, simultaneously with electrical interrogation. This novel neural interface can serve as a universal port to the brain tissue for bi-directional neural recording and stimulation, especially for closed-loop neuromodulation. To implement the Smart Dura, we have devised a microfabrication process that enables parallel, scalable, and high-throughput manufacturing of the devices with customizable arrangements of electrodes.

**Figure 1.**
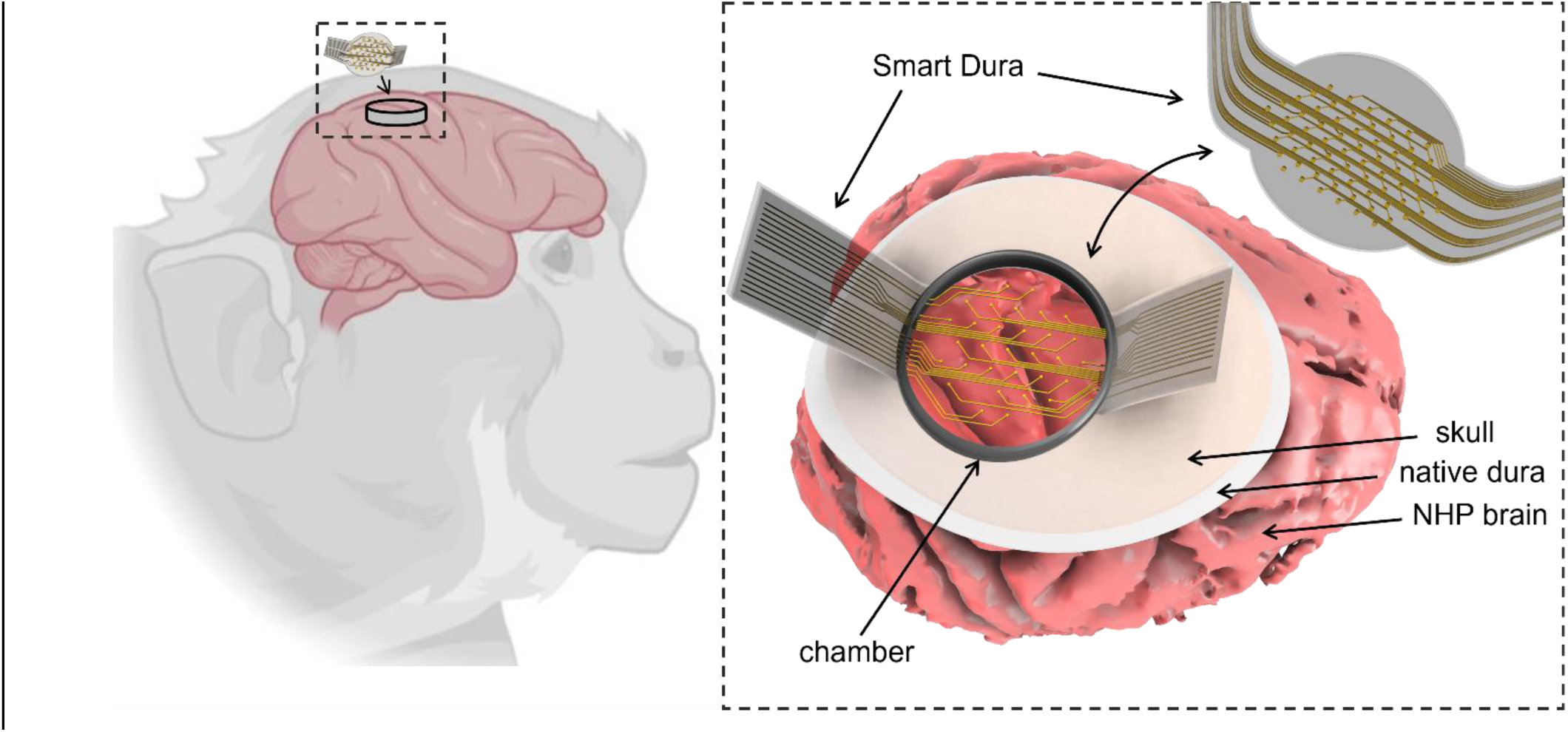
The Smart Dura and its *in vivo* implementation on NHP brain. The Smart Dura is a surface electrode array fabricated on transparent, soft, and flexible polymer substrates that replaces the native dura with a functional device that enables high-resolution distributed electrophysiological recording and electrical stimulation, while providing optical access to the brain for simultaneous multimodal structural and functional optical imaging and neuromodulation.

### Design

The Smart Dura is essentially a surface µECoG array monolithically embedded into a PDMS artificial dura. The Smart Dura provides access to a large area of the brain for recording electrophysiological activity with a high density of recording electrodes. The overall contact surface area of the Smart Dura design in this paper is chosen to be 315 mm^2^ (20 mm in diameter), large enough to cover various areas of the macaque cortex (e.g. motor and somatosensory areas) simultaneously.

We have designed Smart Dura with 32, 64, and 256 electrodes to demonstrate the scalability of our microfabrication process. The electrode pitch of these designs is 2.5, 1.8, and 1.0 mm, respectively. The pitch of each design was selected to maximize the coverage area for electrophysiology, while at the same time, providing the highest level of optical access. Moreover, these designs provide a very high fabrication yield. We also implemented smaller-size devices with 15 mm^2^ (3 mm x 5 mm) area coverage for rodents with electrode pitches of 800, 500, 350, and 200 μm, demonstrating our scalability for high spatial density capabilities with our microfabrication process. It has been shown previously that small electrodes (e.g., 10×10 µm^2^) on surface µECoG arrays can detect single-unit and multi-units in addition to local field potentials (LFPs) from the cortical surface^24^. However, using such small electrodes results in larger noise and deteriorates the signal-to-noise ratio (SNR). Considering this tradeoff, we have designed the electrode sizes in the range of 10-40 µm in diameter (78-1250 µm^2^ surface area) for high spatial resolution recording of electrophysiological signals, while maintaining a high SNR. The high customizability of microfabrication processes in the number of electrodes as well as the electrode size and pitch of our Smart Dura devices allows us to further investigate the effect of these design parameters on resolution as well as spatial and temporal frequency content of electrophysiological signals.

### Substrate material

In contrast to the typical fabrication-centered design of μECoGs, the design of our Smart Dura follows an application-centered approach. The recent trend in μECoG fabrication for increased mechanical compatibility with brain tissue is based on commonly used and commercially available polymer substrates in microfabrication. Thin layers (5-20 μm) of flexible polymers, i.e., polyimide and Parylene C, can provide a high level of flexibility to reduce the mechanical mismatch with brain tissue compared to traditional silicon or other rigid substrates. Despite this improvement, these polymer substrates are still 4 and 3 orders of magnitude stiffer than brain and dura tissue, respectively^25^ (Fig. S1A). To enhance mechanical compatibility, we implemented a thin film bi-layer material stack that forms our Smart Dura using PDMS and Parylene C. PDMS has an elastic modulus (E ∼2 MPa) that is not only 3 times lower than other commonly used polymers, but it is also lower than the elastic modulus of native human dura tissue (E = 60-110 MPa). It has been widely demonstrated that an artificial dura made out of medical grade silicone, such as PDMS, provides an unmatched chronic stability for interfacing with NHP brains^10–12,26,27^. A passive artificial dura made of PDMS can act as a plug or cap to cover open craniotomy and protect brain tissue against infection and unwanted tissue growth between electrophysiology experiments^13,28^. Parylene C has a modulus of elasticity that is three orders of magnitude larger than that of the native human dura. To match the effective modulus of elasticity (i.e., Young’s Modulus) of the native dura, we designed the respective film thicknesses in the material stack to be 250 μm of PDMS and 10 μm of Parylene C to form our Smart Dura (Fig. S1B). This way, not only does our design benefit from the established efficacy and safety of passive artificial dura, but it also provides functional recording and optical access in a unified device that can seamlessly blend with the native dura with a matching effective modulus of elasticity (E = 109.62) to maintain natural cranial pressure.

It should also be noted that performing traditional fabrication processes based on photolithography on PDMS is challenging due to its low surface energy, therefore PDMS is not widely used in microfabricated implants. Our bi-layer substrate design enables us to benefit from the unique mechanical properties and biocompatibility of PDMS, while at the same time, taking advantage of scalable microfabrication processes on Parylene C.

### Fabrication

Our microfabrication process enables the successful incorporation of PDMS into a thin-film material stack for parallel, high-throughput fabrication of metal electrodes and traces (Fig. 2A). Using this microfabrication process, we implemented electrodes as small as 20 μm in diameter, comparable to the size of single neurons. While our processes are capable of 2 µm features, we selected a conservative 10 µm trace width resolution to route the signals from the electrode sites to our back-end connector pads. The electrode and trace dimensions in this work were chosen to address the needs of the intended biology experiments and maximize the optical access, while ensuring a high fabrication yield.

**Figure 2.**
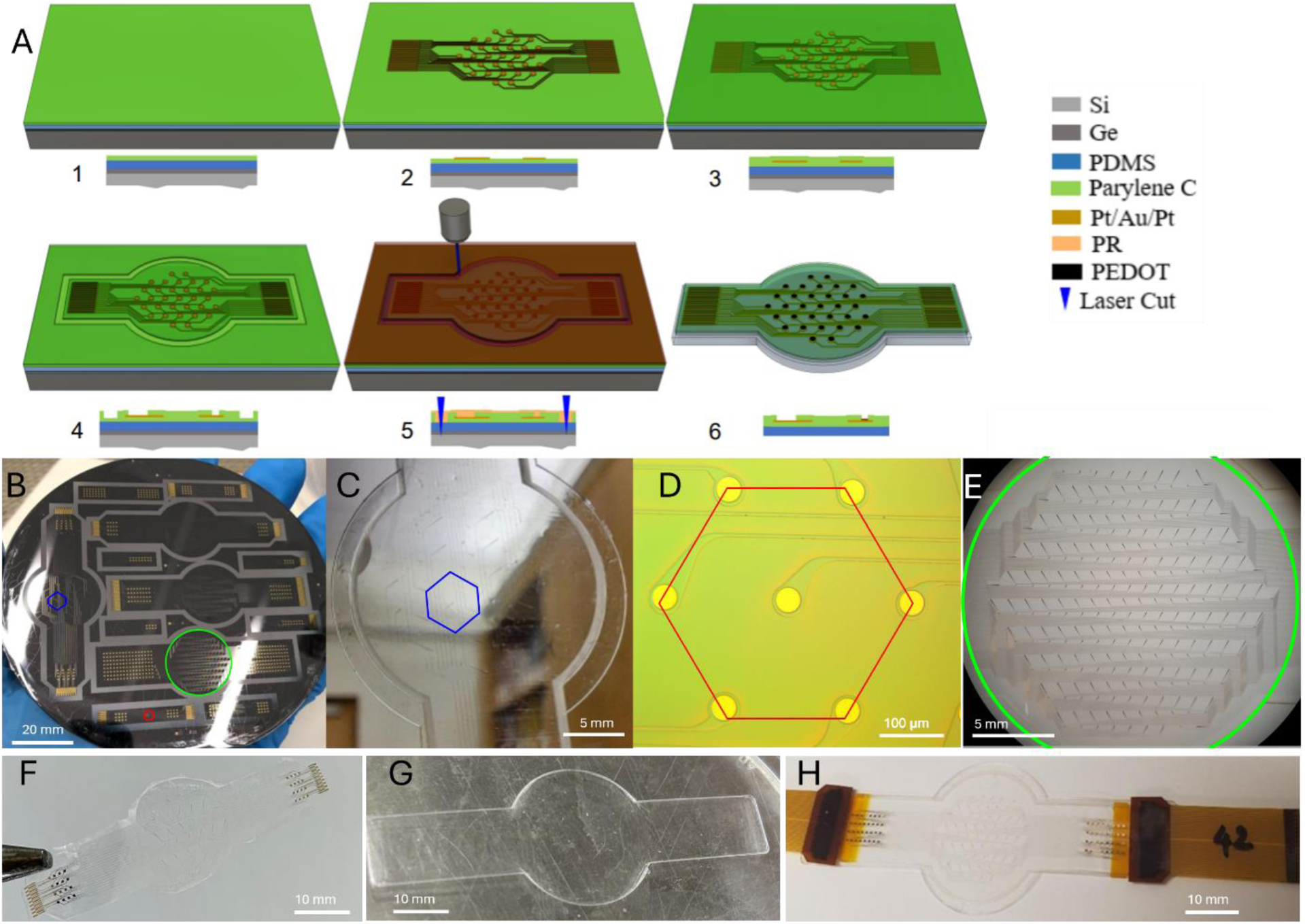
Smart Dura fabrication process flow and features. A) Smart Dura microfabrication process illustrating the layer-by-layer transition in each fabrication step, from deposition of substrates (1), patterning of device features (2-4), to device release (5-6). B) 4-in wafer containing four Smart Dura for NHP application and 8 smaller scale Smart Dura for quality control purposes and small animal models. C) 32-ch Smart Dura with 40 μm diameter electrode and 2.5 mm electrode pitch. D) Micrograph (inset) of Smart Dura with 40 μm diameter electrode and 200 μm electrode pitch array. E) Distribution of the electrodes of our high-density 256-channel Smart Dura over the 20 mm diameter area coverage. F) 32-ch Smart Dura released from the wafer. G) 200 μm thick PDMS backing layer. H) 64-ch Smart Dura packaged with flexible interfacing PCBs for backend connection and with the added PDMS backing layer.

The fabrication of the Smart Dura starts with a bare silicon wafer as the temporary substrate. A thin (100 nm) layer of germanium (Ge) was deposited as the release layer on the surface of the silicon wafer before spin coating PDMS. Once the release layer was deposited, PDMS was spin coated to a 50 µm thick layer (1). Then we deposited a 5 µm layer of parylene C using a chemical vapor deposition method to functionalize the surface of PDMS (1). Parylene C is a biocompatible material widely used as a substrate or insulation layer in implantable neural interfaces, including µECoG arrays^24,29^. It is optically transparent and adheres well to the surface of PDMS when deposited on PDMS. In addition, for thin film devices, Parylene C exhibits lower water vapor transmission rates^30^ compared to PDMS^31^. The following steps followed a planar microfabrication process, previously used for implementing parylene-based ECoGs^24,29^. The recording electrodes and electrical traces were photolithography defined followed by metal deposition and lift-off (2). After the lift-off process, another 5 µm layer of parylene C was conformally deposited as the top electrical insulation layer (3). Parylene C at the electrode sites, contacts, and device outline was etched using a cycled O_2_ plasma reactive ion etching process (3 minutes etch and 2 minutes rest) to expose the electrode sites and the contact pads (4). Once the electrodes and contacts were exposed, the next step was to define the device outlines in PDMS. Although selective dry etching of 50 µm thick PDMS is possible, we resorted to a simpler and quicker laser cutting method to define the outline. Prior to laser cutting, we deposited a thick layer (12 µm) of photoresist to protect the exposed metal electrodes and backend bond pads from the byproducts of laser cutting. PDMS was then removed precisely at the device outline via laser machining (wavelength: 355 nm, kerf: 40 µm) (5). Next, we removed the sacrificial germanium layer by immersing the wafer in a 3% H_2_O_2_ bath for 24 hours and releasing the individual Smart Dura devices (6). The protective layer of photoresist was kept after laser machining to protect the metal electrodes and backend bond pads during the final wet etching step. The protective photoresist is then stripped in acetone, followed by washing steps in isopropanol (IPA) and deionized (DI) water. Hydrogen peroxide (H_2_O_2_) that we used to release the devices is a biocompatible chemical and often used for sterilization. Although we wash the released devices in acetone, IPA, and DI water, any trace amounts of H_2_O_2_ that may remain on the devices would not cause adverse effects due to toxicity. Post fabrication, PEDOT was electro-deposited on the electrodes in a PEDOT:PSS solution to reduce the impedance and improve the SNR^32^.

Exemplary microfabricated NHP Smart Dura devices (32-, 64- and 256-channel designs) are shown in Fig. 2B-H. In each case, we designed a honeycomb electrode lattice for a uniform spacing between electrodes, and edges were filleted to prevent crack formation (Fig. 2D). We packaged the 32- and 64-channel designs directly with zero-insertion-force (ZIF) connectors at the ends. For the 256-channel design, we designed a bond pad array compatible with anisotropic conductive film (ACF) bonding for a more compact packaging. We connected the Smart Dura devices to polyimide flexible PCBs for easier and more robust connectivity to the back-end electronics. Finally, a 200 µm thick backing layer of PDMS that was previously cured via injection molding (Fig. 2G) was attached to the substrate of the device to provide the added PDMS thickness for tunning the effective modulus of elasticity to match the native dura. We attached the backing PDMS layer to the bottom of our Smart Dura with very thin and uncured diluted hexane:PDMS solution (10:1 by weight) that acted as glue. During this process, we coated the ZIF connectors on the flexible PCBs with PDMS to provide mechanical and electrical insulation. Figure 2C and H show the final Smart Dura molded to the backing PDMS layer. The two noticeable edges around the perimeter of the electrode array show the distinction between the edge of the electrode array as released from the wafer and the edge from the backing PDMS layer.

### Electrical and optical characterization

Once the devices were released and packaged, the electrical functionality and optical access were characterized to test the microfabrication and packaging yield, electrical recording and electrical stimulation performance, and the optical access through the Smart Dura devices. The electrochemical impedance of the electrodes in electrolyte mimicking the conductivity of brain tissue was used as a metric to assess the recording performance (baseline noise) and the variation of recording performance from one electrode to another in each device. The charge injection capacity of electrodes was also measured to judge the performance of devices for localized neural stimulation. The optical access through the Smart Dura devices was rigorously measured using optical transmission spectroscopy methods. Along with characterizing the apparent optical transparency of the devices, we present a new framework for defining the overall optical access in the context of multi-photon imaging to demonstrate the microfabricated thin metal traces enable far more optical access beyond the apparent optical transparency.

### Electrochemical impedance spectroscopy

Using a three-electrode electrochemical cell^29^ (Fig. 3A), we measured the electrochemical impedance spectrum (EIS) of all the recording electrodes on each device to assess the electrophysiological recording capability (Fig. 3B). As expected, the impedance magnitude decreased after electrodepositing conductive polymer, PEDOT:PSS (see Methods section). For example, the impedance at 1 kHz, a frequency relevant for single and multi-unit activity, decreased from 971 ± 183 kΩ to 30.4 ± 2.4 kΩ on electrodes with a diameter of 20 µm (sample size, N = 46) (Fig. 3B) and from 365 ± 46.8 kΩ to 13.4 ± 0.48 kΩ for 40 µm diameter electrodes (sample size, N = 32). At 100 Hz, a frequency more relevant for LFP signals, the impedance decreased from 6.39 ± 1.19 MΩ to 64.7 ± 2.7 kΩ on electrodes with a diameter of 20 µm (Fig. 3B) and from 2.34 ± 0.30 MΩ to 29.6 ± 1.25 kΩ for 40 µm diameter electrodes. The small standard deviation of impedances at all measured frequencies shows that the variation of impedance is less than 10% across all the electrodes for 20 µm electrodes and less than 5% for 40 µm. Given such low variation of impedance across electrodes and since thermal noise is proportional to the square root of the real part of impedance^33^, the variation of the noise performance is further reduced. Based on the average impedance values we measured, the thermal noise^33^ at 30 kHz (sampling rate of our electrophysiology system) of our 20 µm and 40 µm diameter electrodes is 3.382 µV_rms_ and 2.264 µV_rms_, respectively. Hence, the noise floor of our Smart Dura devices is low and comparable to the noise floor of our electrophysiology recording equipment and consistent across all electrodes. For LFP, multi-unit and single-unit signal levels in the typical range of 150 µV to 1mV, our SNR would be at a minimum of 75 dB, considering a 150 µV signal and our 20 µm diameter 3.382 µV_rms_ noise level. The yield of the devices, defined as the number of working channels divided by the total number of electrodes, was 100% for most of our 32-channel devices. The yield was reduced to around 96% for higher channel count devices. The limited yield was mainly due to broken traces during the lift-off process or handling issues during the packaging steps.

**Figure 3.**
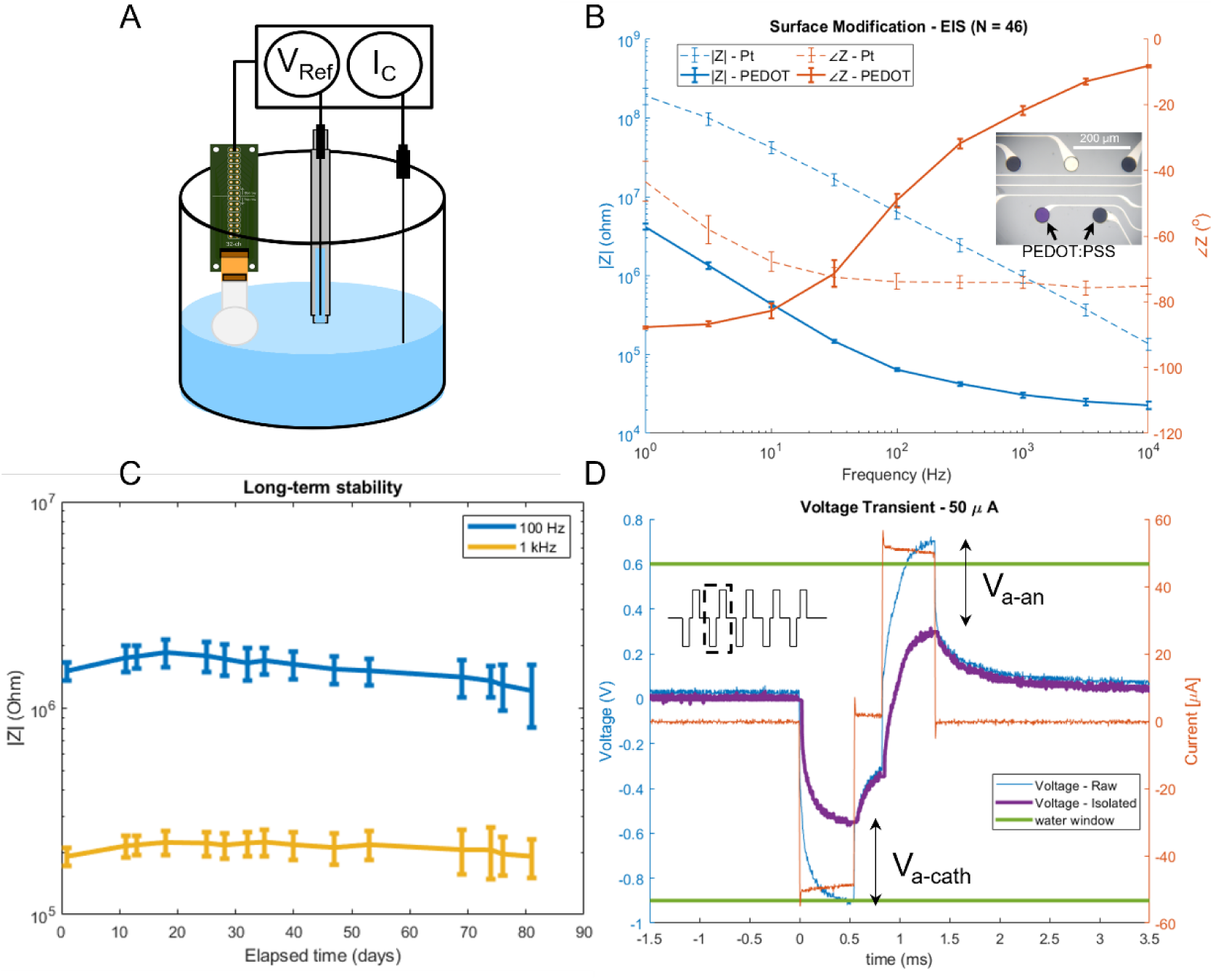
Electrical characterization of Smart Dura. A) Electrochemical cell with a three-electrode configuration (counter, reference, and working electrode) used for the electrical characterization of our Smart Dura. B) EIS measurements of a 20 µm diameter electrode with the native Pt surface and PEDOT:PSS surface modification. Inset micrograph of electrodes with native Pt electrodes and PEDOT:PSS coated electrodes. C) Long-term stability test of our Smart Dura immersed in 1x PBS solution for 81 days. Average and standard deviation of the electrochemical impedance magnitude vs time of 40 µm diameter Pt electrodes at 1 kHz and 100 Hz shows the stable long-term performance of our Smart Dura without degradation. D) Voltage transient measurements response to a burst of bi-phasic current waveforms of PEDOT:PSS electrodes. The blue trace shows the raw voltage measurement and the purple trace shows the isolated voltage transient waveform across the electrode after removing the near-instantaneous voltage artifact (V_a-cath_ and V_a-an_) at the onset or termination of the current pulse (see Fig. S3).

### Long-term stability

In order to determine the long-term stability of our Smart Dura to remain functional for periods of time that exceed those of acute applications, we performed a lifetime stability test that consisted in immersing our Smart Dura in 1x PBS solution continuously at room temperature for an extended period of time. The PBS solution contains the main chemical ions that are present in the brain and dominate its electrical conductivity, making this solution appropriate for stability testing. EIS measurements were taken periodically to obtain electrode impedance data during the extended period of time. As seen in figure 3C, the average and standard deviation of the electrochemical impedance magnitude vs time of 40 µm diameter Pt electrodes at 1 kHz and 100 Hz remains consistent over time and within the two standard deviation window. These results show the stable long-term performance of our Smart Dura without degradation for up to 81 days.

### Charge-balanced stimulation

Using the same three-electrode electrochemical cell arrangement as in EIS, we injected a train of charge-balanced current pulses to characterize the stimulation performance of our Smart Dura devices by calculating current injection capacity (CIC). As detailed in the Methods section, to determine the CIC we repeated the injection of charge-balanced current pulses with a frequency and pulse width tailored for our NHP experiments^34^ with increasing current amplitudes until the measured voltage levels exceeded safe limits (i.e. the water window). Operating within the safe electrochemical window avoids irreversible reactions and the production of toxic compounds such as H^+^ and OH^-^ from the electrolysis of water in the tissue. For PEDOT:PSS-coated electrodes used in the Smart Dura, the water window is −0.9 to 0.6 V^35^.

As shown in Fig. 3D, the current amplitude achieved with our Smart Dura before reaching the water window (E_mc_ of −0.48 V and E_ma_ of 0.31 V) was 50 μA. Using the procedure explained in the methods section, the CIC for our electrodes coated with PEDOT was measured to be 8.754 mC/cm^2^, commensurate with the maximum amount reported for PEDOT:PSS^35^ and ∼20 times greater than an equivalent Pt electrode of the same electrode size.

### Optical access

One of the key advantages of Smart Dura is that it provides optical access for simultaneous optical imaging and electrophysiological recording. The optical access through Smart Dura is enabled by using a transparent substrate as well as microfabricated traces and electrodes. To characterize the optical properties of Smart Dura, we measured the direct optical transmission using the setup shown in Fig. 4A. A broadband light source from an incandescent light bulb illuminated the sample from the bottom via a 45-degree broadband aluminum mirror. A fiber-coupled spectrometer (200 µm core diameter, Ocean Optics HR4000CG-UV-VIS) was used to measure the transmission spectrum. The fiber was placed directly above the Smart Dura surface. We first measured the optical transmission of Smart Dura through air. This is how such biomedical devices are usually characterized for optical transparency. However, in practice, Smart Dura will be interfacing with the brain tissue on the transmission side. To mimic this realistic condition, we performed measurements with de-ionized (DI) water added on top of Smart Dura using a PDMS chamber (Fig. 4A). The fiber tip was then immersed in water to characterize the optical transmission through the Smart Dura. In this measurement, DI water is used since its refractive index approximates the average background refractive index in real tissue^36^. As shown in Fig. 4C, the optical transmission through the Smart Dura substrate is greater than 80% for the visible spectrum (400-750 nm) and 85% for the near-infrared (780-1000 nm) regions, which are relevant wavelength ranges for neurophotonics methods such as optogenetics and calcium imaging. The optical transmission spectrum is dominated by the PDMS layer and is almost identical to the PDMS transmission spectrum alone^37^. Parylene C exhibits very little absorption in the visible and near-infrared range^30^. The metal stack used in the traces and the electrodes render those regions opaque. The surface area blocked by the metal traces and electrodes is only 1.58% of the total 32-channel Smart Dura surface area. Therefore, 98.42% of the Smart Dura surface (32-channel design) area exhibits larger than 80% transmission in the visible and infrared range of the spectrum. However, the levels of transparency of the Smart Dura are improved when implanted on the surface of the brain, in contact with the cerebral spinal fluid (CSF). CSF has the same refractive index as DI water (*n* = 1.33), which exhibits less contrast with the substrate (i.e., Parylene C with *n* = 1.64), compared to air (n = 1). The refractive index of the backing PDMS layer is *n* = 1.40. The reflection coefficient at each of these interfaces can be analytically calculated. The reflection at the Parylene C and air interface (5.88%) is significantly higher than the reflection at the Parylene C and tissue (DI water) interface (1.09%), a 4.79% difference, as shown in Fig 4B. Figure 4C shows the transmission spectra through the air (black curve) and through air into the water (blue curve). Figure 4D shows the difference between the two spectra, which demonstrates a more than 2% improvement across the entire visible spectrum when DI water was added on top of the Smart Dura surface. The discrepancy between the theoretical prediction of improvement compared to the experimental results is partly because our simplified analysis only considers light reflection effects at the input and output interfaces, neglecting multiple partial reflections that result in Fabry Perot etalon effects^38^. Moreover, light attenuation due to material absorption and scattering from the surface is not considered. The decrease in transmission in the infrared regions (>800 nm) is due to the dominant effect of water absorption^39^.

**Figure 4.**
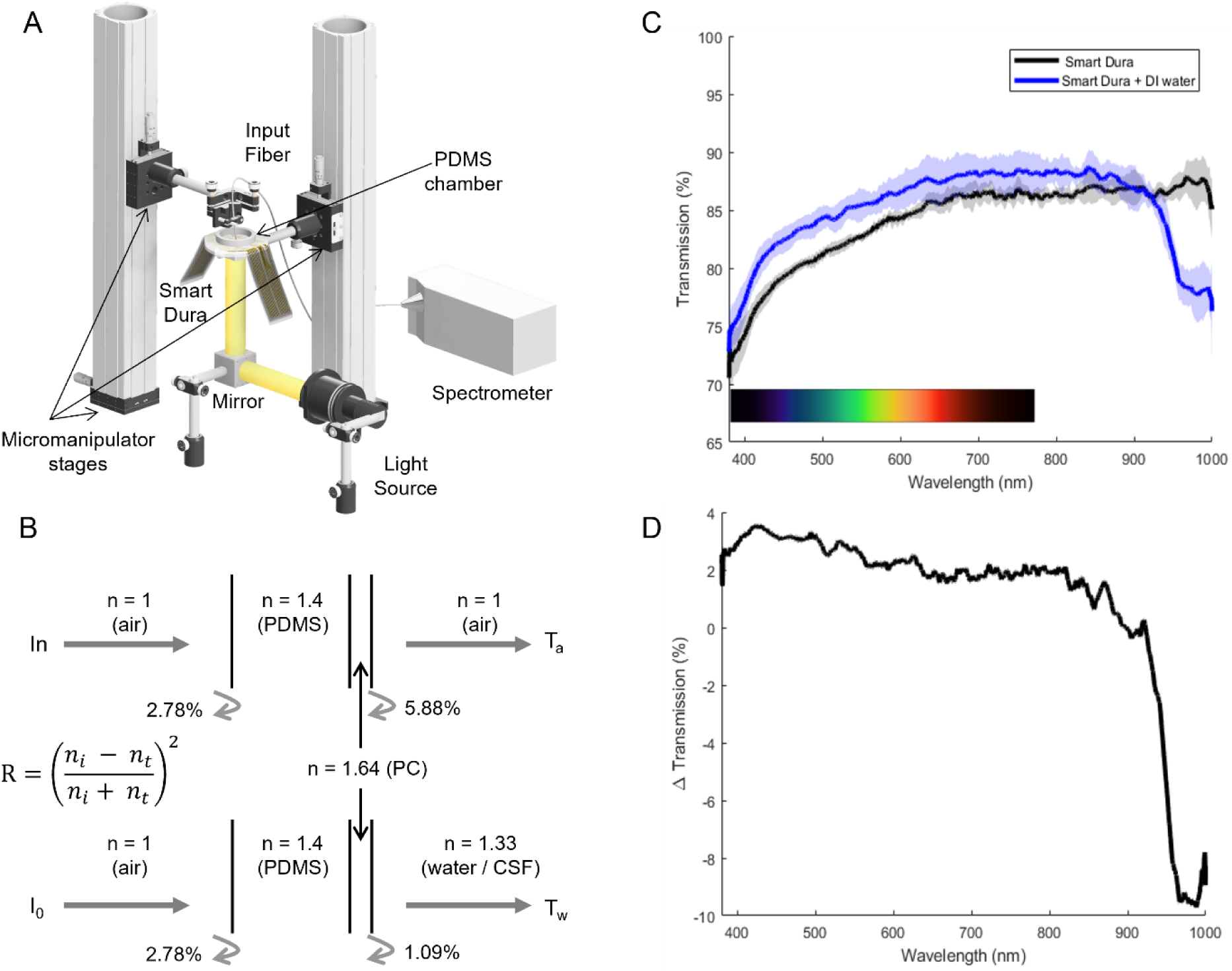
Optical transmission characterization of Smart Dura. A) Schematic of the optical setup for transmission spectra analysis. B) Analytical calculations of light reflected at input and output material interface of the Smart Dura and air (top) and Smart Dura and water (bottom), showing the difference in light reflected between Parylene C and air vs Parylene C and water. C) Direct transmission spectrum of the Smart Dura alone (black trace) and Smart Dura and water (blue trace). D) 2% transmission improvement for the visible range with the added water interface vs the Smart Dura alone. These results demonstrate a high level of transparency with a transmission level of >80% across the visible spectrum of the Smart Dura alone. Additionally, in our application, we benefit from the CSF layer interface as it provides a better-matched transition of light transmission into the brain.

It should also be noted that under *in vivo* conditions, we usually image the structure or function of neurons or vasculature at a certain depth below the surface of the brain, i.e., the target imaging plane is not the Smart Dura surface. In this case, the optical access depends on how the rays of light can pass through the Smart Dura to reach the target imaging depth. With the surface electrode arrays on Smart Dura, we can record the collective electrophysiological activity of a population of neurons across several cortical layers. Simultaneous functional optical recording using calcium indicators such as GCaMP will provide complementary insights into the cellular activity of individual neurons. In particular, multi-photon imaging allows deep optical access by focusing the optical beam at the target depth. Since the image plane is typically a sub-millimeter to a millimeter below the surface of the brain where the cell bodies of cortical neurons are located, the Smart Dura will not be at the focal plane and therefore, if the traces are narrow enough, light can be focused around the traces.

Figure 5A shows the Smart Dura placed on the cortex (sensorimotor cortex of monkey M). As shown, the transparency of the Smart Dura provides optical access to the underlying tissue, as evidenced by the high-resolution image captured with a digital camera (FinePix XP70; FUJIFILM Co.), revealing the underlying microvasculature and cortical areas. We conducted two-photon fluorescent imaging of the vasculature through the Smart Dura in the cortex (primary visual cortex (V1) of monkey N). As shown in Fig. 5B, immediately after injecting FITC-dextran intravenously, we were able to get clear images of microvasculature under the Smart Dura. Despite the traces being opaque, as shown in Fig. 5B and C, light can pass around the narrow metal traces that form interconnects (ROI2, Norm. intensity: 0.978), but the wider parts and the larger electrode area with a diameter of 50 µm cast shadows on the optical images (ROI1, Norm. intensity: 0.403). This finding shows the benefit of using lithographic techniques to fabricate the Smart Dura and achieve trace and electrode geometries in the micron scale, which become practically transparent for optical methods where the light source or imaging field of view extends to regions much larger than the size of the electrodes and traces. This makes the Smart Dura optically transparent (with a transmission >80%) for neuroscientific methods and shows its potential to be combined with optical stimulation or imaging techniques.

**Figure 5.**
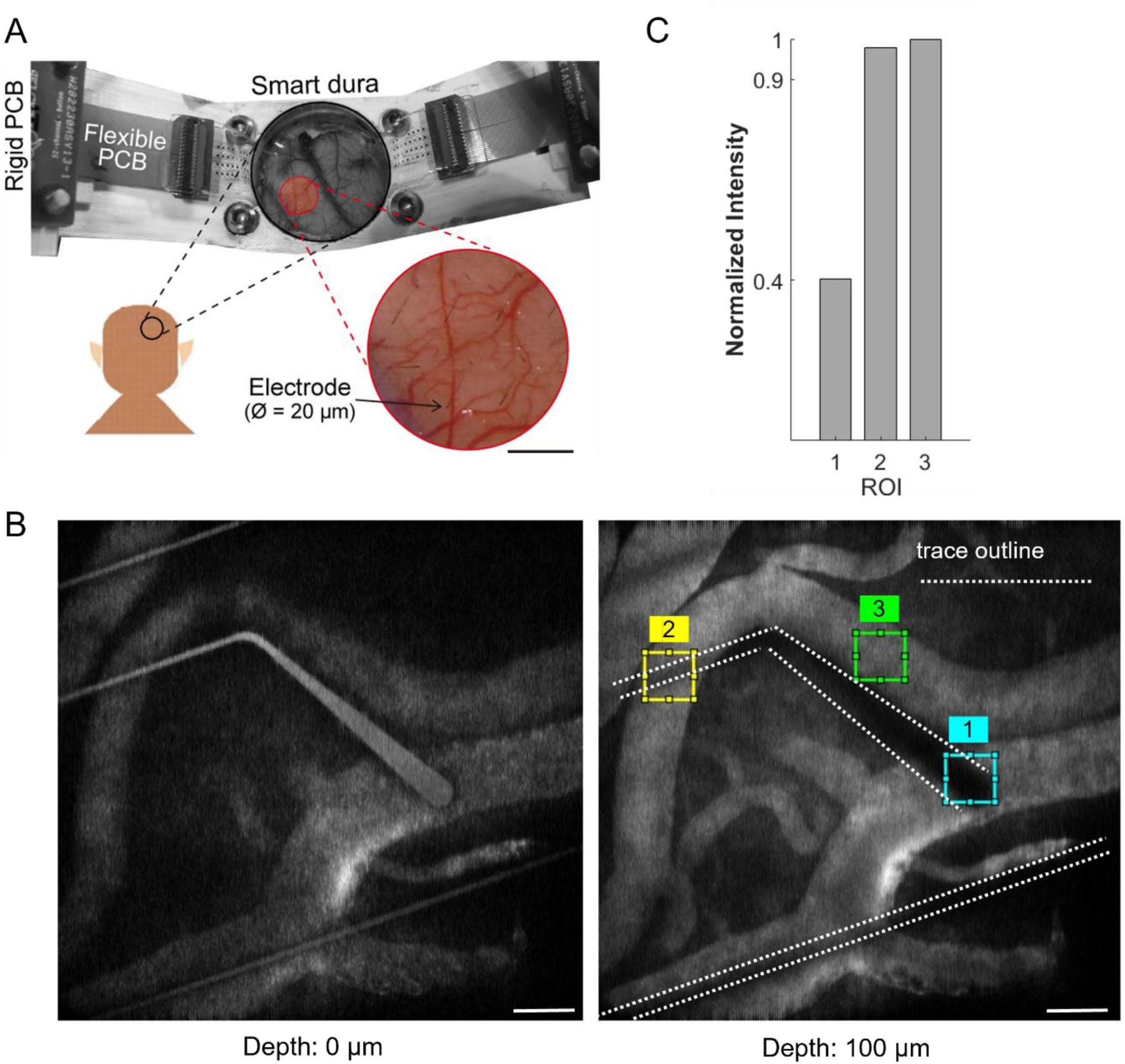
In vivo optical characterization of Smart Dura with two-photon microscopy. A) Photograph showing the microelectrode placement over the cortical surface. Scale bar: 2 mm. B) *In vivo* fluorescent vascular images in the primary visual cortex through the Smart Dura using two-photon microscopy, focused on the metal traces of the Smart Dura (depth: 0 µm) and the vasculature below (depth: 100 µm) with three regions of interest (ROI) of size 44×44 µm^2^ over the electrode site, where the metal measures 50 µm in diameter (1), around a 10 µm trace (2) and an unobstructed area (3). Scale bar: 100 µm. C) Bar plot comparing the average greyscale values of the three regions of interest, showing how a 10 µm trace is practically transparent as no appreciable signal is lost compared to the electrode site.

Table 1 summarizes the electrical and optical characterization results.

**Table 1.**
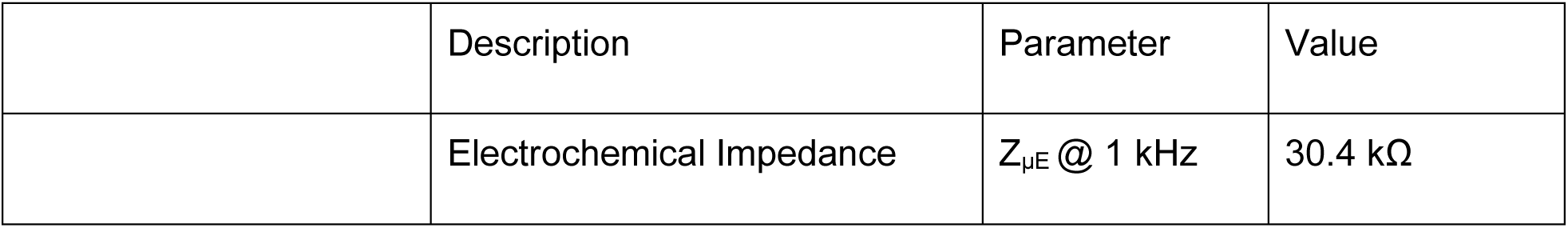
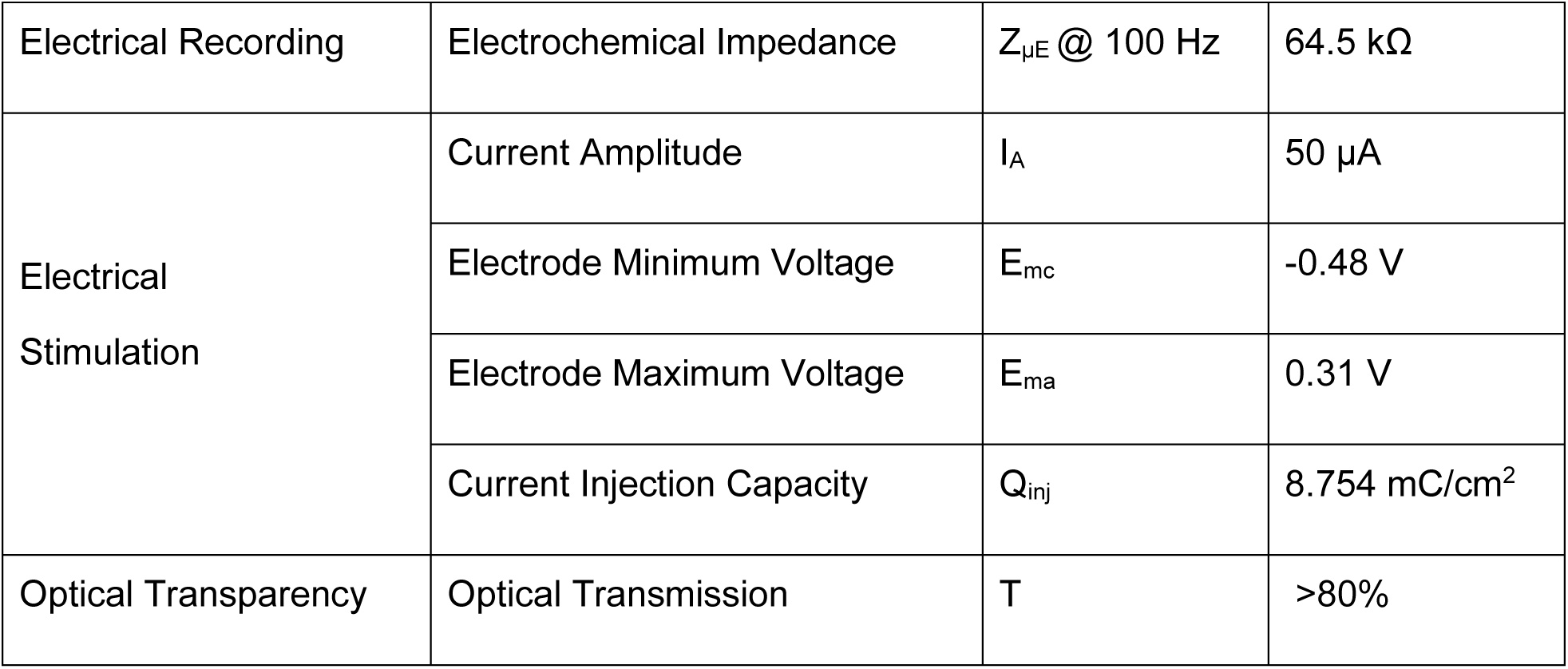
Electrical and optical properties of Smart Dura. The metrics associated with the electrode characterization correspond to the electrode with a 20 µm diameter.

|  | Description | Parameter | Value |
| --- | --- | --- | --- |
| | Electrochemical Impedance | $Z_{\mu E}$ @ 1 kHz | 30.4 kΩ |
| Electrical Recording | Electrochemical Impedance | $Z_{\mu E} @ 100 \text{ Hz}$ | 64.5 kΩ |
| Electrical Stimulation | Current Amplitude | $I_A$ | 50 $\mu\text{A}$ |
| | Electrode Minimum Voltage | $E_{mc}$ | -0.48 V |
| | Electrode Maximum Voltage | $E_{ma}$ | 0.31 V |
| | Current Injection Capacity | $Q_{inj}$ | 8.754 mC/cm<sup>2</sup> |
| Optical Transparency | Optical Transmission | T | >80% |

### *In vivo* neural recording in NHP brains

Our Smart Dura is a multi-modal interface designed for use in various experimental settings to understand brain function and dysfunction, develop and evaluate therapies for neurological and neuropsychiatric disorders, and combine electrical and optical techniques with behavioral analysis to unravel the connection between brain networks and functions. To demonstrate and validate the versatility of the Smart Dura for these applications, we conducted *in vivo* tests in both anesthetized and awake animals, during behavioral tasks and at rest, as well as following stimulation, across multiple cortical areas.

### Electrophysiological neural recording from NHPs

As described before, the Smart Dura was designed with electrode diameters optimized to record highly localized high-frequency activity in NHP cortices. To evaluate this capability under different conditions, we recorded neural signals from two macaques in both anesthetized and awake states, which are known to differ in level and complexity of neural activity and responsiveness to external stimuli, and from two cortical areas (V1 of anesthetized monkey N and posterior parietal cortex (PPC) of awake monkey H). We recorded electrical signals at a sampling rate of 30 kHz and extracted LFP and spiking activity by filtering the signal into low- and high-frequency bands with a cutoff frequency of 250 Hz (left panels in Fig. 6A, B). From the high-frequency band signals, we isolated single- and multi-units in both monkeys (right panels in Fig. 6A, B). From the waveforms obtained, we computed our empirical SNR (using equation 7 in the methods section) of the spikes detected to be 45 dB for spike in figure 6A and 32 dB for spike in figure 6B. The difference between the theoretical SNR values from the EIS measurements (reported earlier in the electrical characterization section) compared to the empirically measured

**Figure 6.**
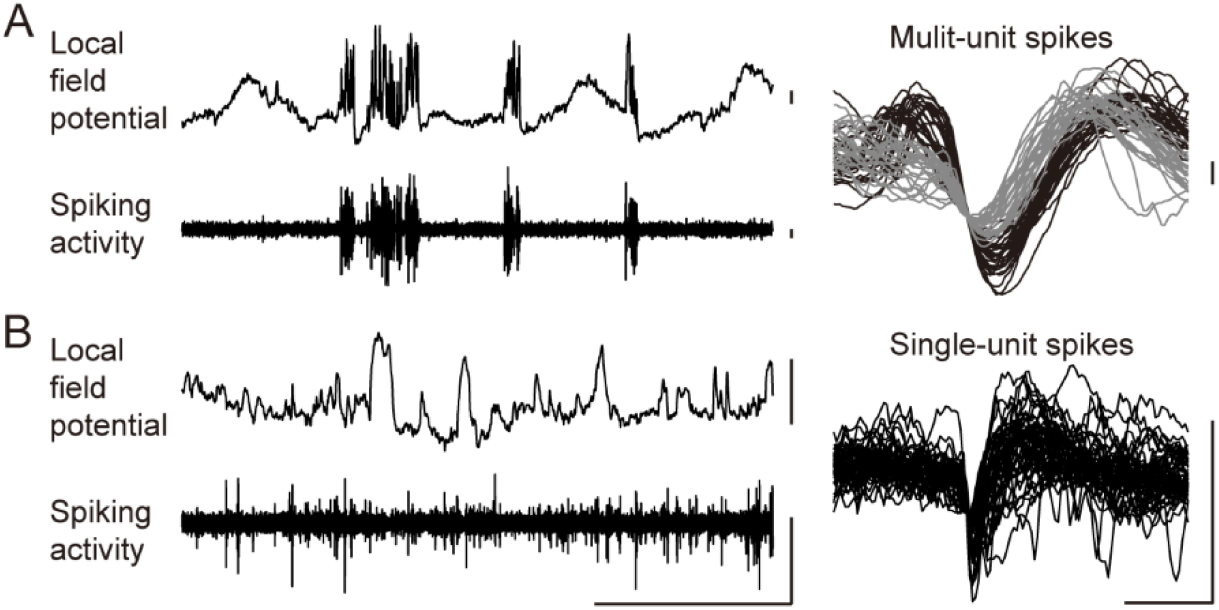
Electrophysiological recording using the Smart Dura in NHP cortex. LFP (frequency band: < 250 Hz), spiking activity (frequency band: > 250 Hz), and spike waveforms of isolated units recorded from a single electrode of the Smart Dura in the visual cortex of anesthetized monkey N (A) or in the posterior parietal cortex of awake monkey H (B). Waveforms of different colors represent different sorted units. Scale bars for amplitude: 1 mV, 200 μV, and 150 μV for LFP, spiking activity, and single-unit spikes, respectively. Scale bars for time: 1 s and 1 ms for signal traces and spike waveforms, respectively.

SNR values is largely due to the additional sources of noise present during the *in vivo* experimental procedure that are not accounted for in the theoretical SNR calculations. The additional sources of noise are the 60-Hz (and harmonics) power line noise, or spurious electromagnetic noise from equipment used for the completion of the experiment such as electrically driven pumps or actuators. Nevertheless, 45 and 32 dB empirical SNR values are within the same range of SNR values obtained from intracortical neural interfaces in NHP^40^. These results demonstrate the capability to record not only LFPs but single-and multi-unit activity from the surface of the NHP brain using our Smart Dura.

### Neurophysiological recording in correlation with behavioral tasks

The Smart Dura was tested in an awake-behaving animal to capture task-related neural activity during behavioral tasks. Initially, we recorded epidural activity from the left PPC of monkey H using both a 32 and 64-channel Smart Dura in separate experiments while the animal performed an instructed-delay center-out reaching task (Fig. 7). Analysis of the epidural signal power between the task and resting periods was conducted for each electrode and averaged across trials, with both arrays containing at least 800 successful trials included for this experimental condition. We observed a significant increase in the low-frequency theta (4-8 Hz) power in multiple electrodes for both arrays (8 of the 32-electrode array and in 6 of the 64-electrode array) when comparing neural activity during behavior with rest (first row in Fig. 7A). This activity was spatially localized and consistent across both arrays, with enhanced spatial resolution evident in the 64-channel array with its higher electrode density (2500 µm and 1818 µm electrode pitch for 32- and 64-channel arrays, respectively). Examples of LFP power histograms are shown to compare the power changes during the task in both array types. No significant increase in theta power was observed in the control trials taken from randomly shuffled resting data (second row in Fig. 7A), suggesting that these recorded changes were specific to the task and its associated neural processes. Subsequently, similar recordings were obtained subdurally using a 64-channel Smart Dura. Notably, we detected a significant increase in low-frequency theta power (4-8 Hz) similar to the epidural setup (first row in Fig. 7B). Moreover, this subdural approach revealed significant enhancements in task-related higher frequency gamma power (30-70 Hz), which were not observed in the epidural recordings (first row in Fig. 7B). This finding suggests that subdural recordings, being closer to the neuronal sources, capture a broader spectrum of frequency content, including higher frequencies that appear attenuated or absent in epidural recordings.

**Figure 7.**
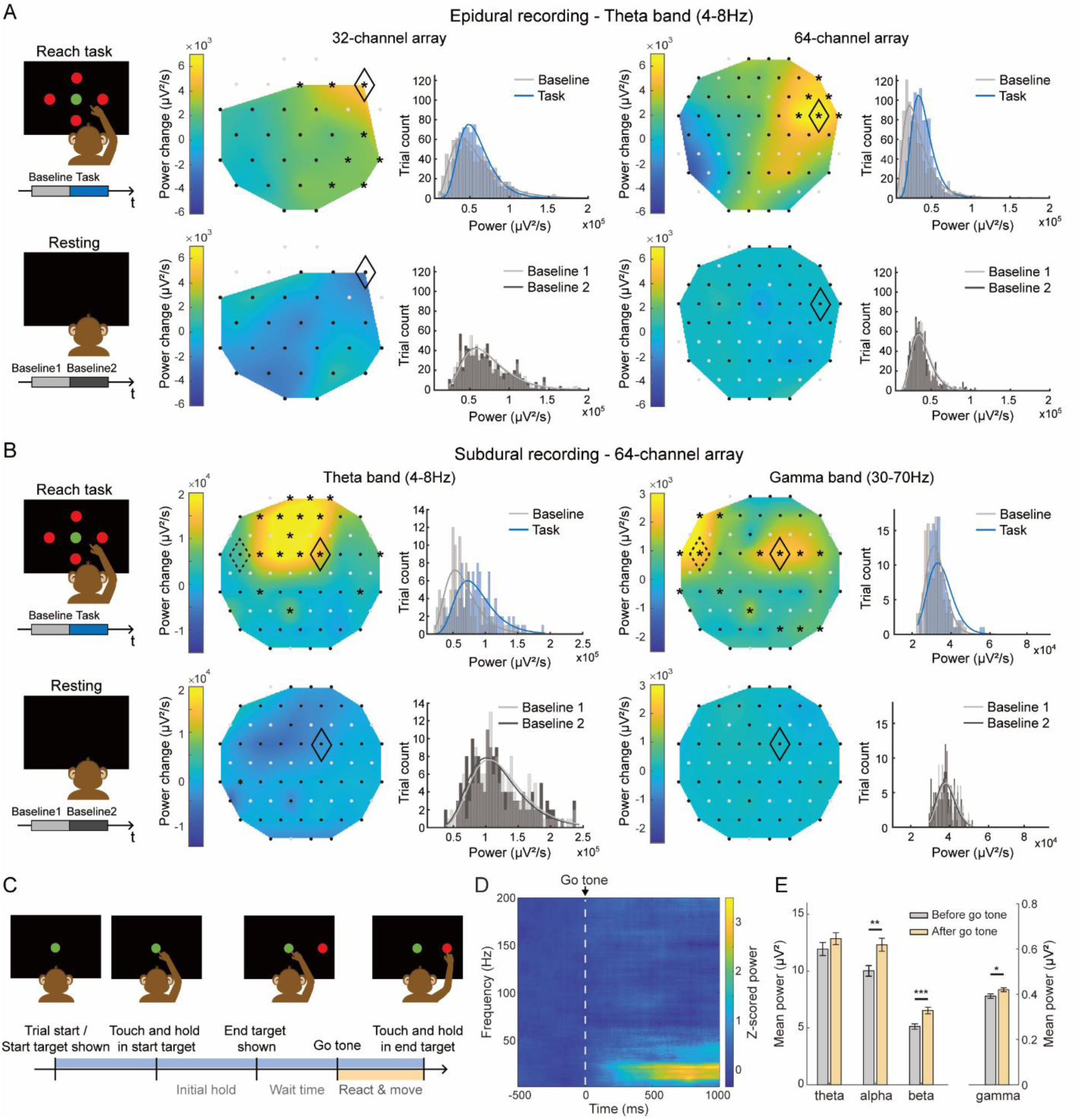
Large-scale recording in the posterior parietal cortex during a center-out reaching task. A) Epidural recording and signal power in the theta band (4-8 Hz) during the reach task (first row) and resting periods (second row), collected using a 32-channel (middle) and a 64-channel array (right), respectively. Heatmap showing the trial-averaged power change between task and baseline segments or during resting periods. The locations of the recording electrodes and the electrodes with significant power change were labeled in black dots and asterisks, respectively. Histogram showing the signal power distribution across trials for the selected electrode, marked with a black diamond on each heatmap. B) Subdural recording and signal power analysis with the 64-channel array during the reach task (first row) and resting periods (second row) in the theta (4-8 Hz; middle panel) and gamma (30-70 Hz; right panel) frequency bands. Black dashed diamonds in the heatmap denote the electrode used to generate the subsequent spectrogram. C) Schematics showing the detailed timeline of each trial during the center-out reaching task. D) Spectrogram showing the trial-averaged time-frequency distribution of subdural LFP signals during the reach task from 500 ms before to 1000 ms after the go-tone. Spectrogram values were z-scored to the segment before the go-tone. E) Bar plots of the averaged LFP power at individual frequency bands (theta, alpha, beta, and gamma) based on the before go-tone and after go-tone recording segments averaged across 162 trials (*p<0.05, **p<0.01, ***p<0.001; paired t-test).

Next, we characterized the movement-specific neural response by realigning the subdural LFP recordings to go-tone (Fig. 7C). The *z*-scored spectrogram revealed that a high response electrode (black dashed diamond in Fig. 7B), middle to higher frequency bands were selectively increased after the go-tone (Fig. 7D). Bar plots also revealed this trend (Fig. 7E), showing that the average power after go-tone was significantly higher than before go-tone in the alpha, beta, and gamma frequency bands (paired t-test, p<0.05). Together, these results suggest that the Smart Dura is capable of capturing location and frequency-specific neural response both during the entire task period and during the reach movement alone. More broadly, our ability to record detailed neural activity in both epidural and subdural modalities highlights the utility for studying cortical dynamics across brain regions. Given the role of the posterior parietal cortex in integrating sensory information and coordinating motor actions^41^, the Smart Dura provides a valuable tool for investigating neural processes underlying diverse cognitive and motor functions.

### Neural recording of evoked responses to sensory and electrical stimulation

Neural interfaces that allow neuromodulation with simultaneous recording of evoked responses are critical for neuroscience research. To demonstrate this functionality of our Smart Dura, we used two different stimuli as representatives: sensory stimulation and electrical stimulation. First, to show that the Smart Dura can detect sensory-evoked neural activity in response to peripheral tactile stimulation, we recorded LFPs using a 64-channel Smart Dura from the right hemisphere of the sensorimotor cortex in anesthetized monkey M while stimulating the left-hand fingertips using a custom-made vibrating stimulator (Fig. 8A, B). As shown in Fig. 8C, applying stimulation to all fingers resulted in a simultaneously evoked response in the somatosensory cortex. The sensory-evoked response in the low-frequency band (4-15 Hz) was statistically significant in the most lateral region in the array (2 electrodes marked with black asterisks in Fig. 8C; p = 0.014, p = 0.0097). This lateral somatosensory response is known to represent finger sensation in humans and macaques^42^. In the significant electrode (D in Fig. 8C), the power increase after stimulation onset was notable below 15 Hz, which was most profoundly observed during the first 200 ms of stimulation, as seen in the spectrogram in Fig. 8D. Such vibrotactile-evoked activity changes in low-frequency bands have been reported in previous studies in human subjects using ECoG or EEG^43,44^. These results show that our Smart Dura can be used to obtain sensory-evoked information from the somatosensory cortex.

**Figure 8.**
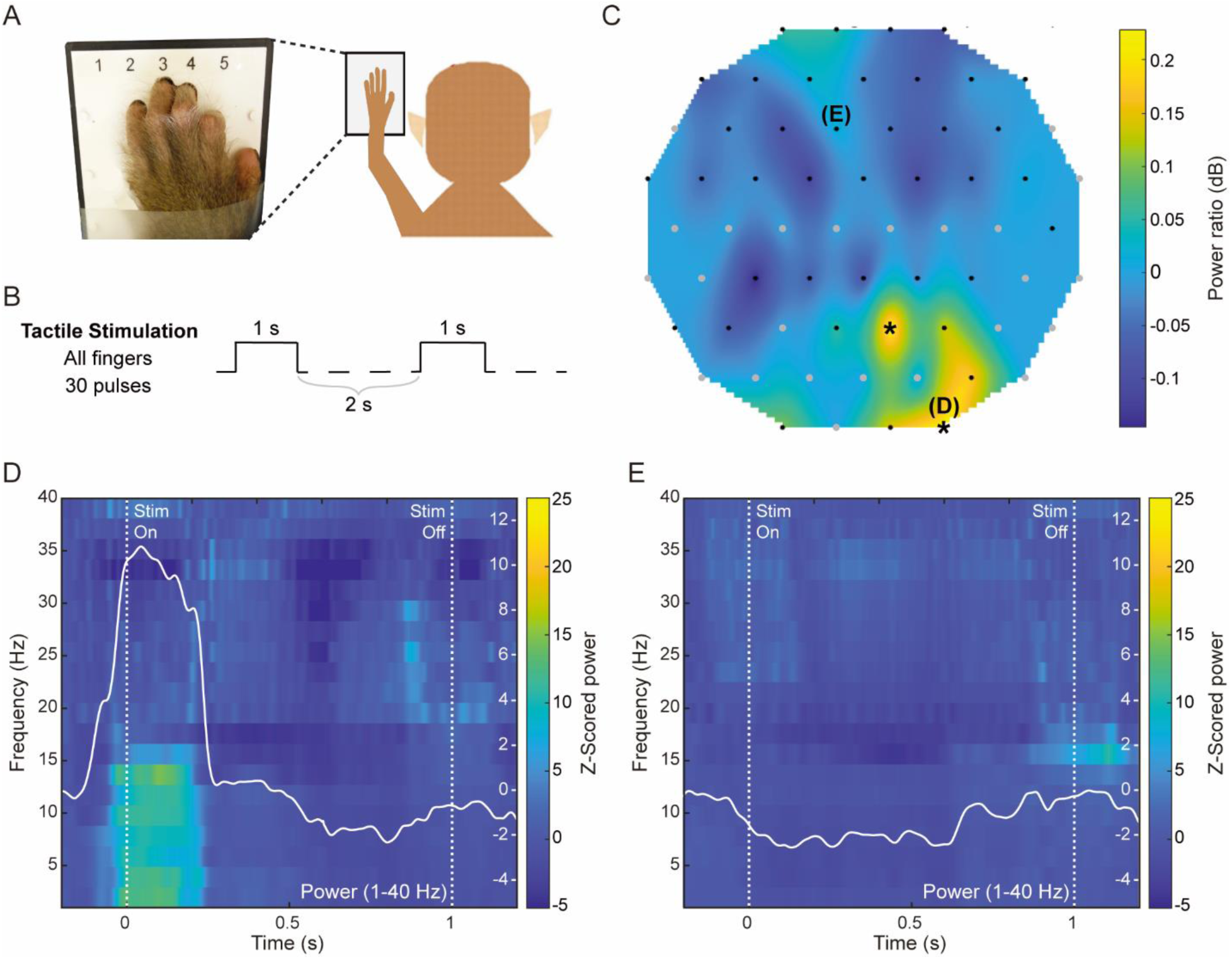
Locating evoked sensory activity in the somatosensory cortex. A) Experimental setup in which monkey M’s fingers are placed over individual tactile stimulators. B) Tactile stimulation pulses. C) Heatmap of signal power change in the low-frequency band (4-15 Hz) during tactile stimulation of all five fingers simultaneously. Black dots show recording electrodes, and black asterisks denote the electrodes with significantly increasing power during stimulation (p<0.05; two-sample t-test). D-E) Spectrograms of individual electrodes showing a sensory-evoked response (D in Fig. 8C) and no evoked response (E in Fig. 8C) during tactile stimulation. The stimulation pulse onset occurred at t = 0 and lasted 1 second. For each spectrogram, the averaged power in the 1-40 Hz frequency band is overlaid in white.

Next, to assess the ability of Smart Dura to record and stimulate neural activity, we recorded LFPs epidurally in the PPC of awake monkey H with a 64-channel Smart Dura and electrically stimulated with a paired stimulation paradigm. One block within a recording session consisted of two consecutive baseline 10-minute periods, followed by a 10-minute period of paired electrical stimulation of two cortical sites, then a 10-minute period of spontaneous post-stimulation recording (Fig. 9A). We analyzed low (30-59 Hz) and high gamma (60-150 Hz) band power to assess local changes in neural activity throughout the underlying cortex. To measure the network effects of epidural stimulation, we compared spontaneous and stimulation-induced changes by calculating the change in power between both initial baseline periods and the post-stimulation and the initial baseline period for each block. As a result of stimulation, we observed increases in regions near and distant to the stimulation sites in comparison with spontaneous activity changes (Fig. 9B). Across the network, there was a significant increase in network-wide activity levels following stimulation in both frequency bands compared to the spontaneous activity changes (Fig. 9C). These stimulation-induced changes demonstrate the ability to electrically stimulate and record cortical activity epidurally with the Smart Dura.

**Figure 9.**
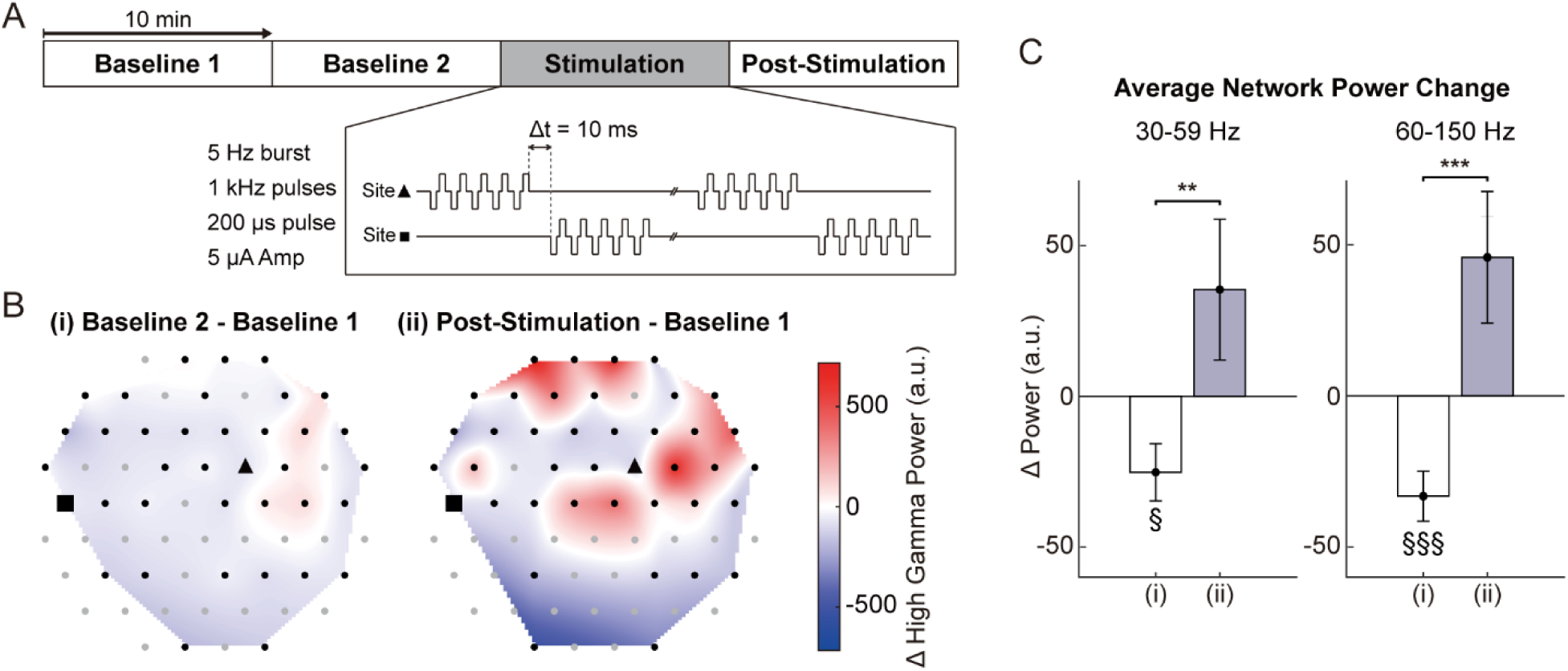
Alteration of neural network activity evoked by epidural electrical stimulation. A) Time course for each block of epidural recording and electrical stimulation. Two baseline periods of spontaneous LFPs were recorded, lasting 10 minutes, followed by a 10-minute period of paired epidural electrical stimulation of two cortical sites. A fourth 10-minute period was used to record post-stimulation spontaneous activity. B) Heatmap of mean high gamma (60-150 Hz) power change for an example block. The left heat map demonstrates spontaneous changes in activity levels, while the right heat map shows stimulation-induced activity changes across the array. Black dots show recording electrodes. A triangle and a rectangle denote the stimulation electrodes. C) Comparison of spontaneous and stimulation-induced changes in average network low (30-59 Hz) and high gamma (60-150 Hz) band power. One-sample t-tests were used to test for changes in power relative to the Baseline 1 period (low gamma spontaneous changes: p = 0.0082; low gamma stimulation-induced changes: p = 0.13; high gamma spontaneous changes: p = 7.26×10-5; high gamma stimulation-induced changes: p = 0.036). Paired sample t-tests were used to compare across groups (low gamma: p = 0.0011; high gamma p = 2.16×10-5).

### Optogenetic stimulation by local light illumination

As discussed earlier, Smart Dura provides a very high level of optical access to the brain tissue underneath the array. This enables a plethora of multimodal optical neuromodulation and recording techniques, simultaneously with electrophysiology recording and electrical stimulation. To demonstrate the multimodality of Smart Dura, we applied optogenetic stimulation to the PPC of monkey H that had opsin expression of a red-light-activated Jaws. We placed a 64-channel Smart Dura and positioned an optical fiber coupled with a red-light laser at the wavelength of 638 nm between two electrodes (the red star in Fig. 10A) located within the region, where opsin was expressed (White dashed lines in Fig. 10A). We then delivered light stimuli with a laser power of 45 mW while recording the evoked neural response. Figure 10A shows the heatmap of the power change during the stimulation that was z-scored with respect to the power in the pre-stimulation period and divided by the impedances of individual electrodes to eliminate the influence of impedance differences between electrodes. During light stimulation, we observed reliable light-evoked neural activity near the stimulation site (Fig. 10A) that was absent in the baseline recording without stimulation (Fig. 10B). We quantified the power changes at four electrodes located at different distances from the optical stimulation zone. As shown in Fig. 10C, significant power increases were observed only at the electrode closest to the stimulation zone. These results demonstrate that the Smart Dura can be applied for targeted optogenetic modulation while simultaneously monitoring neural activity. Additionally, these results showcase the potential of our Smart Dura to perform closed-loop neural interfacing with high spatial and temporal resolution neural stimulation via optogenetics and network level electrophysiology recording of LFP activity from the surface of the brain over large regions of the brain. These capabilities demonstrate the true multimodal ability of our Smart Dura for neural interfacing of NHP brain.

**Figure 10.**
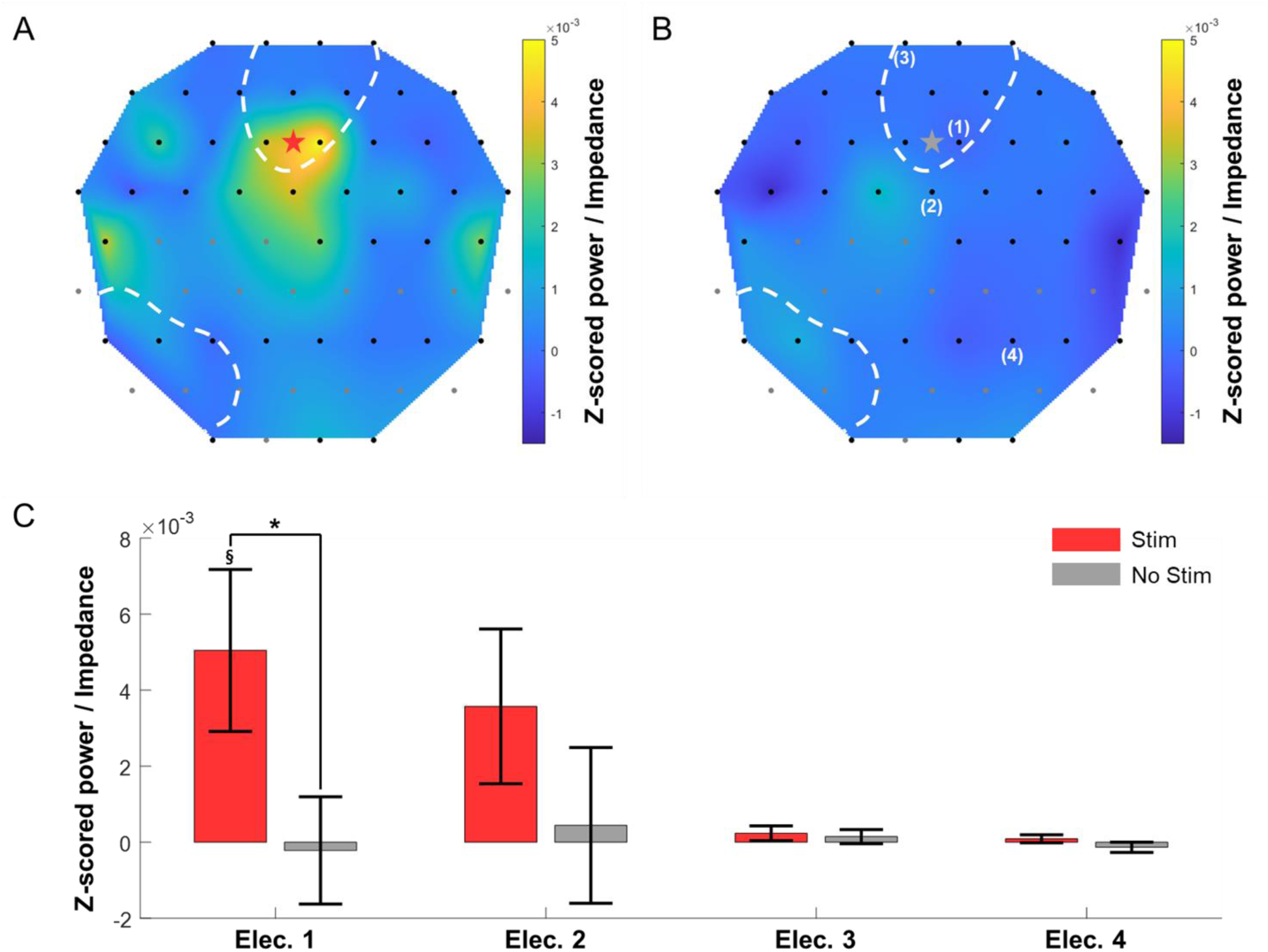
Targeted optogenetic stimulation and evoked neural response. A) Heatmap of power change in the low-frequency band (4-40 Hz) during light stimulation with a laser power of 45 mW. White dashed lines and a red star show opsin expression areas and laser location, respectively. Black dots denote recording electrodes. B) Heatmap of power change in the low-frequency band (4-40 Hz) during baseline recording without laser stimulation. C) Power change at four different electrodes, shown in Fig. 10B. Data are presented as the mean ± standard error of the mean. One-sample t-test to test for changes in power relative to the pre-stimulation period (§p = 0.0247). Two-sample t-test to compare with baseline recording without stimulation (*p = 0.0454). n = 30 trials.

## Discussion

In this paper, we demonstrated the Smart Dura, a microfabricated customizable multimodal surface neural interface with mechanical properties similar to the native dura and therefore, can seamlessly blend with the native dura for high-resolution electrophysiology and targeted localized electrical neural stimulation. The careful choice of substrate materials to provide a high level of optical transmission, along with miniaturized microfabricated metal traces and electrodes, provides a very high level of optical access for multimodal optical imaging and optogenetic neuromodulation. Our scalable microfabrication and packaging process enables high throughput implementation of Smart Dura with customizable arrangement and size of electrodes.

### Material choice and tunable mechanical properties

The implementation of our Smart Dura can be modeled by the simple case of a circular membrane fixed at its edges with a uniform pressure load applied at the bottom of the membrane. Given the large area coverage of 3.14 cm2 (20 mm diameter) of our Smart Dura, the mechanical interaction between the native tissue and our Smart Dura is described by the flexural rigidity of the device. The flexural rigidity of our bi-layer membrane depends on the geometrical parameters (i.e., the diameter and thicknesses) and the bulk material properties (i.e., the Young’s Moduli and Poisson ratio) of the combined individual layers. As shown in figure S1A, the modulus of elasticity of PDMS is 3 orders of magnitude softer than Parylene C and 1 order of magnitude softer than native dura mater, while Parylene C is 2 orders of magnitude stiffer than native tissue. We chose our PDMS and Parylene C layers to be 250 µm and 10 µm, respectively, tune the effective stiffness of our Smart Dura to be on the softer range between PDMS and Parylene C levels and closer to the mechanical properties of native dura tissue. The simulation results are shown in figure S1B-E. We used finite element method (FEM) solid mechanics simulations using COMSOL Multiphysics. As shown in the cross-section view in figure S1C, our 250 µm and 10 µm PDMS/Parylene C bi-layer substrate exhibits a maximum deformation at the center of the membrane of 1.825 mm. Comparing these results to a different thickness configuration of 220 µm and 40 µm PDMS/Parylene C bi-layer substrate we can observe the notably smaller deformation (0.304 mm) and hence a stiffer performance due to the larger thickness ratio of Parylene C. These results show how the effective stiffness of a bi-layer PDMS + Parylene C substrate can be tuned in our design by choosing the relative ratio of the Parylene C and PDMS thicknesses. The capability of tunning the effective mechanical properties our Smart Dura enabled by our bi-layer microfabrication process highlights the versatility in our design for enhanced mechanical compliance with native brain and dura tissue.

### Optical access vs high density electrodes

Compared to the current state-of-the-art functional artificial dura, i.e., the MMAD^15^, our Smart Dura provides significant improvements in scalability and optical access (Fig. S2). Both improvements are primarily driven by the increased spatial resolution of our microfabrication process. Nanoparticle inkjet printing-based technology used in the MMAD has a fabrication resolution of 200 µm trace width, which limited the electrode count to an upper bound of 32 electrodes per circular area of 20 mm in diameter, or 315 mm^2^ (Fig. S2A). The MMAD achieves an increase in optical access by stacking four 8-electrode µECoG films complementarily designed on top of each other to decrease the number of light-blocking traces. However, with the miniaturization of the minimum feature of our Smart Dura down to 10 µm trace width, we could significantly improve the optical access and scalability (Fig. S2C and D). For a one-to-one comparison, we calculated the area of the optical window not blocked by the opaque metal traces and electrodes in the MMAD and Smart Dura, respectively (Fig. S2E). Compared to the optical access of 87.84% from the 32-channel MMAD, our 32-channel Smart Dura provides an optical access of 98.42%, rendering it very transparent. More notably, our 256-channel Smart Dura still provides higher optical access than the MMAD (90.68%) while providing eight times the number of electrodes. This shows the powerful improvement of our Smart Dura performance compared to the MMAD based on the increased minimum size feature resolution of our microfabrication process.

Furthermore, following the trend from Fig. S2E, one can argue that by doubling the channel count from 256 to 512 electrodes, the optical access will be reduced by at least another 10%. This will decrease the optical access (to 80% or below), compromising the performance for multi-modal applications and setting an upper limit on the electrode count based on the desired optical access. However, our microfabrication process, which can implement thin metal traces on the order of 10 µm, enables further enhanced transparency. As shown in Fig. 5B and C, despite the metal traces being opaque, we could still image vasculatures underneath the 10 µm-wide traces. This is possible when traces are separated far enough (> 20 µm), like in our Smart Dura designs, to allow enough light to go around the traces and be captured by the imaging system, rendering the individual traces effectively transparent. Performing the same calculations for the optical window area but neglecting the 10 µm width traces, the optical access of our Smart Dura increased to 99.64%, 99.46%, and 98.69% for the 32, 64, and 256-channel designs, respectively (Fig. S2E). Therefore, the high-resolution microfabricated traces provide a unique opportunity to achieve effective optical access that is way beyond the apparent optical transparency, especially when the traces are routed such that densely packed concentrated areas are avoided in the Smart Dura design.

Effectively, only the transition tapers from the traces to the electrodes, and the electrodes themselves block the optical access of our Smart Dura devices. To achieve such a high level of effective optical access, the size of traces and the tapers can be even further reduced using higher-resolution microfabrication. The resolution limit of our fabrication process is driven by factors affecting the planarization of our material stack and the image reversal resist (AZ-5214) we use during metal patterning via lift-off. Given the film thickness of our AZ-5214 resist of 1.4 µm, and by optimizing our material stack planarization, the resolution limit of our microfabrication process can be further scaled from 10 µm down to 2 µm. Therefore, instead of fabricating 10 µm wide traces, we could reduce the size down to 2 µm trace width to obtain practically transparent traces again for at least 5 times the electrode count and density than demonstrated in this paper for the same circular area coverage of 20 mm in diameter, or 315 mm^2^. In this case, an electrode array of 1024 electrodes would still provide optical access higher than 90%, at 93.45%. Moreover, in future iterations, the larger taper and electrode geometries with 20-50 µm diameter in size could be made transparent by implementing alternative transparent-conductive materials such as Indium Tin Oxide (ITO) or graphene in the microfabrication process flow to further increase the optical access.

### Multi-modal neural interfacing

The very high level of optical access through our Smart Dura enables simultaneous electrophysiological recordings with optical imaging and neuromodulation techniques for multi-modal interfacing with the brain. For example, the 96% optical access of our 256-channel Smart Dura, enables high-density electrophysiology simultaneously with a plethora of optical methods such as functional multiphoton calcium imaging to acquire neural activity at single-cell resolution in the deeper brain layers of the cortex. Combining this with the neural recording capabilities of the Smart Dura, which provides large-scale surface recordings with high temporal resolution, can open up new opportunities to integrate complementary information about the underlying physiology. In addition, the photothrombotic technique^45^ with optical coherence tomography angiography or wide-field hemodynamic imaging, which can induce focal ischemic lesions and image vascular dynamics, can be conjugated with our electrophysiological recordings to further understand neurovascular functions.

### Single- and multi-unit activity recording

The micron-scale electrodes of our Smart Dura are capable of recording single- and multi-unit activity (Fig. 6) as well as LFPs (Figs. 7, 8, 9) from the cortical surface. Surface recording using an ECoG or μECoG array generally measures network-wide signals of neuronal populations. In particular, high-frequency LFPs, such as the high gamma band, reflect responses of more spatially localized areas and highly correlate with spiking activity^46^. However, it is still challenging to pick up spikes of individual neurons. There have been few studies that have succeeded in capturing the spiking activity of individual neurons using surface μECoG arrays with similar features to ours with an electrode size of 20 µm. Recently, a μECoG array with electrode sizes of 10×10 µm^2^ or 10 µm^2^ was reported to acquire spikes from behaving rats and human patients with epilepsy^24,47^. Another recent study reported μECoG arrays with 20 µm diameter electrodes to record spiking activity in songbirds^48^. However, to our knowledge, our paper is the first report of neural recording of single- and multi-unit activity from superficial cortical layers in NHPs. The ability of our Smart Dura to record both LFPs and spikes facilitates obtaining richer neural information in a less invasive manner for NHP studies.

### Customizability and multi-scale recording

Our Smart Dura can potentially enable long-term, large-scale multimodal/bidirectional interfacing with the NHP brain. Leveraging the customizability of our microfabrication techniques together with a co-designed chamber, the Smart Dura provides a platform for multi-scale recording of cortical activity, where the electrode array specifications can be configured for the desired application. The number, size, and density of the electrodes can be selected from a range of designs and be swapped in place in the chamber based on the needs of the study. For example, when interested in the large-scale network connectivity over the whole area under investigation (i.e. M1 and S1) a Smart Dura with ‘large’ (e.g., 50 um) electrodes and distributed over the entire region can be used to record the LFP activity of the large populations of neurons. Additionally, if it is desired to localize the activity of a specific region, the Smart Dura can be swapped by an equivalent one with a higher density of electrodes and with smaller electrode sizes, for recording the LFP and multi-unit and single-unit activity of smaller populations of neurons. We believe that developing such an advanced multi-modal interface in translational NHP models will provide significant insights and opportunities for the development of bi-directional therapies for neurological disorders, especially in closed-loop therapeutical paradigms.

## Conclusion

We have developed a novel multi-modal neural interface, named Smart Dura, which is a functional version of the artificial dura with integrated electrophysiological electrodes for large cortical area coverage for the NHP brain. The Smart Dura is fabricated using a planar micromachining process on PDMS and Parylene C to monolithically integrate a micron-scale electrode array into a soft, flexible, and transparent substrate to provide matched mechanical compliance with the native tissue while providing high-density electrodes (up to 256 electrodes) and achieving high optical access (exceeding 97%). Our *in vivo* experiments demonstrate electrophysiology capabilities combined with neuromodulation, as well as optical transparency via multiphoton imaging. This novel neural interface is an important addition to the neuroscientific toolset for large-scale, bidirectional interfacing for multimodal and closed-loop neuromodulation capabilities to study cortical brain activity in non-human primates, which provides great translational potential to humans.

## Methods

### Smart Dura Fabrication

Building on the design of parylene-based µECoGs^29^, the Smart Dura fabrication process includes additional initial steps for the implementation of PDMS in the material stack and as the device substrate. The complete fabrication process, illustrated in Fig 2A, was performed as follows:

A silicon (Si) wafer was cleaned in acetone, isopropanol (IPA), and water to start the process on a clean and particle-free surface. Before depositing our substrate material (PDMS), germanium (Ge) was thermally evaporated (Angstrom Covap II, Angstrom Engineering Inc. Ontario, Canada) for a 100 nm release layer Fig. 2A(1). Next, PDMS (Sylgard 184, Dow Corning Corp, USA) was spin-coated for a 50 µm thick layer substrate Fig. 2A (1). As mentioned above, we chose PDMS as the material for the Smart Dura because of its biocompatibility, elasticity, and optical transparency. However, PDMS is difficult to integrate into traditional fabrication processes. Because of its low surface energy, it cannot be wetted with photoresist. To address this issue, we deposited a 5 µm layer of Parylene C via chemical vapor deposition (Labcoter 2, Specialty Coating Systems, Indianapolis, IN) to functionalize the surface Fig. 2A (1). Parylene C is optically transparent and adheres well to the surface of PDMS. The next step was to define the recording electrodes and electrical traces. The Pt/Au/Pt metal stack provides good adhesion to Parylene C and at the same time has high enough electrical conductivity. To deposit the metal stack, a 1.5 μm thick image reversal (AZ5214E) photoresist was patterned using contact lithography (MA6 contact aligner, Karl Süss, Garching, Germany). Following the development in AZ400K (MicroChemicals), the wafer surface was cleaned in IPA and water, and the exposed parylene C was activated using O_2_ plasma in Trion RIE (Phantom RIE, Trion, Clearwater, FL) at 100 W and 100 mTorr for 30 seconds. Pt (10 nm)/Au (100 nm)/Pt (10 nm) was deposited on the patterned photoresist by electron-beam evaporation (Kurt J. Lesker PVD 75 Electron Beam Evaporator, Jefferson Hills PA). Excess metal was lifted off in an acetone bath for two hours plus successive rinses were done in IPA and DI water for 5 minutes Fig. 2A(2).

After the lift-off process and prior to the deposition of a second Parylene C layer, wafers were cleaned, and the parylene C was activated in O_2_ plasma at 100 W and 100 mTorr for 30 seconds and then dehydrated at 95 °C for 30 minutes. A 5 µm layer of parylene C was conformally deposited to provide electrical insulation Fig. 2A(3). An aluminum hard mask was deposited via electron-beam evaporation and patterned via wet etching ahead of the electrode and contact opening. Parylene C at the electrode sites and contacts was exposed with a cycled O_2_ reactive ion etch: 3 minutes etch (50 W, 50 mTorr, O_2_: 60 sccm) and 2 minutes rest. A total of 6 cycles (24 min etch) was needed for the removal of the 5 µm layer at the desired locations. After etching, the etched trenches were measured using a surface profilometer (P-15 Stylus Profiler, KLA Tencor, USA), to confirm the electrodes and connector pads were exposed Fig. 2A(4). Next, a thick layer (12 µm) of photoresist (AZ4620) was deposited ahead of laser cutting and device release for protection to laser residue and H_2_O_2_ during device release. Then, the device outline was precisely cut using a UV laser cutter (LPKF ProtoLaser U4, Germany) with 5.7 W maximum laser power at 355 nm Fig. 2A(5). Finally, the wafer was immersed in a 3% H_2_O_2_ bath for 24 hours to dissolve the Ge and release the Smart Dura devices. Post fabrication, electrode surface modification was performed to deposit PEDOT:PSS Fig. 2A(6) via electroplating (Autolab, PGSTAT204, Metrohm, Switzerland) to reduce the electrochemical impedance^32^.

### Electrical Impedance Spectroscopy

For the EIS measurements, we prepared a phosphate-buffered-saline (PBS) solution with a similar conductivity to the cerebrospinal fluid (CSF) in the brain^49^. We used a three-electrode configuration with an Ag/Ag-Cl reference electrode, the electrodes on the Smart Dura as the working electrodes, and a tungsten counter electrode with a large surface area compared to the Smart Dura electrodes, as shown in (Fig. 3A). The reference voltage from the Ag/Ag-Cl electrode and the current through the tungsten electrode were used to measure the electrochemical impedance through the electrode-electrolyte interface (Eq. 1). A modular potentiostat/galvanostat machine (Autolab – Metrohm Co, Switzerland) was used for these measurements.

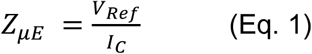

The electrochemical impedance of electrodes in neural interfaces is a standard metric used in the field to categorize the level of transduction of the electrode to convert the electrochemical signal in the extracellular space to an electric signal used in the recording equipment. Both the magnitude and phase of the impedance provide insightful information on the electrical behavior of the electrode-electrolyte interface with respect to signal frequency. Knowing the electrochemical impedance of the electrodes is important in determining whether the electrodes are compatible with electrophysiological recording and stimulation systems, or moreover the working parameters such as amplification or filtering, among others, for optimal recording and stimulation of neural activity. Additionally, the electrochemical impedance values obtained through equation 1 provides a baseline signal to noise ratio (SNR) level of our electrodes by considering its intrinsic thermal noise. The theoretical SNR of our electrodes can be computed from equation 2 below.

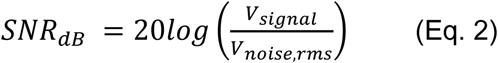

The noise term in the denominator is obtained from the thermal noise model of a resistive device provided by equation 3 below.

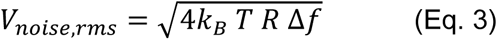

where 𝑘_𝐵_ is Boltzmann constant (1.38*10-23 J/K), *T* is temperature (300 K), *R* is the real part of our electrode (R = |*Z*| cos𝛳, where |*Z*| and 𝛳 are the impedance amplitude and phase) and Δ𝑓 is the frequency bandwidth taken here at 30 kHz given the sampling frequency of our recording equipment of 30 kilo samples per second.

### Electroplating of PEDOT

Electroplating PEDOT on Pt electrodes was performed to reduce the electrochemical impedance of the Smart Dura electrodes by ∼30 fold. To achieve this, we prepared an EDOT:PSS (3,4-Ethylenedioxythiophene:Poly(sodium 4-styrenesulfonate)) solution with a concentration of 0.01 M and 5 M of EDOT and PSS respectively on DI water. For our purposes 80 mL of the EDOT:PSS solution was enough to electrodeposit PEDOT on all the electrodes for any of our Smart Dura devices. Starting with 80 mL of DI water, we added 96 mg of PSS and mixed it with a magnetic stirrer at room temperature for one hour. Then 80 µL of EDOT monomer was mixed into the solution and stirred for an additional hour. With this EDOT:PSS solution, we electrodeposited PEDOT onto the electrodes using the same three-electrode chemical cell shown in Fig. 3A and potentiostat equipment as for EIS. A voltage of 0.8 V was applied to each electrode to polymerize EDOT into PEDOT on top of each electrode. To ensure complete polymer film coverage on the surface and obtain a uniform coating on each electrode, we tracked the amount of charge accumulated during electrodeposition and set a cut-off limit of 1µC/µm^2^. Hence, for our Smart Dura electrodes with 20 and 40 µm diameter sizes, the cut-off charge limit was 315 µC and 1256 µC respectively. Figure 3C shows a comparison between a Smart Dura array before and after PEDOT deposition on certain electrodes selected. There is a noticeable difference in the color of the electrodes between the native Pt electrodes and the PEDOT:PSS deposited electrodes. The characteristic black color of PEDOT:PSS coated electrodes is due to the rougher surface which disperses the incident light in different directions and results in little or no light reflected back to the microscope objective^50^.

### Charge-balanced Stimulation

In addition to the electrochemical impedance, the electrical stimulation capabilities of electrodes are characterized by determining the voltage levels across the electrode-electrolyte interface during the current stimulation waveforms intended for stimulation. Voltage transient measurements *in vitro* are used to estimate the maximum amount of current that can be injected within safe voltage bounds. In studies or therapies involving electrical neural stimulation, the current waveforms typically used are charged-balanced current pulses to reverse all electrochemical reactions back to chemical equilibrium^35^. Depending on the application (DBS, epilepsy, etc.), the shape of the current pulses can vary. For electrical stimulation from the surface of the brain on NHPs, it is typical to use a burst of 5-10 bi-phasic current pulses with pulse widths of 200-1000 μs and shorter (50-200 μs) interphase periods^34^. The reason for this pulse topology with burst is to increase the effectiveness of the stimulation over a large population of neurons.

As in EIS, we used the same three-electrode electrochemical cell configuration where the current pulse is injected with the counter electrode and through the working electrode, and the voltage transient at the electrolyte is recorded by the reference electrode. A burst of 5 bi-phasic charged-balanced current pulses with a pulse-width of 0.6 ms and an interphase period of 200 μs was used for these measurements. The amplitude of the current pulse (I_A_) was increased starting from 1 μA in a cyclical manner until the maximum voltage levels within safe bounds were determined. It is important to note here that the voltage transient recorded at the electrolyte with the reference electrode is not exclusively the voltage drop across the electrode-electrolyte interface. The initial electrode polarization or bias level (E_ipp_) and the electrolyte’s resistance, overpotential, and concentration gradient (usually included in the activation potential term, V_a_, for both anodic and cathodic pulses) also contribute to the voltage recorded. To determine accurate metrics, the voltage across the electrode-electrolyte interface must be isolated from the raw measurements. Once isolated, the figures of merit used to characterize electrodes, such as Charge Injection Capacity (Q_inj_), the voltage across electrode-electrolyte interface (V_μE_), and their minimum and maximum values (E_mc_ and E_ma_) can be obtained, as illustrated in Fig. S3. With the voltage transients’ raw measurements, the following process, illustrated in Fig. S3, was performed to extract the figure of merit values:

1. Calculate the electrode initial polarization (E_ipp_) and activation potential V_a_ for the cathodic and anodic pulses to isolate the voltage across the electrode-electrolyte interface (V_μE_) following the signal processing steps described by Cisnal et al^51^.
2. To calculate E_ipp_, V_a-cath_, and V_a-an_, first take the 1st derivative of the voltage recorded to find t_Va-cath_ using the finite central difference expression

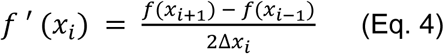

i. b. With t_Va-cath_, calculate E_ipp_ by taking the average of baseline value before the cathodic pulse:

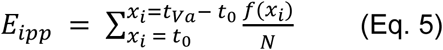

i. c. Then subtract E_ipp_ from the voltage transient to remove this offset term.
ii. d. Lastly, determine the activation potential, V_a-cath_, and V_a-an_ and adjust the waveform to remove the V_a-cath_, and V_a-an_ contributions to obtain the isolated voltage across the electrode-electrolyte (V_μE_).
iii. 2) With V_μE_, obtain E_mc_ (min value) and E_ma_ (max value).
iv. 3) From the geometrical surface area (GSA) of the electrode, the current amplitude, and the pulse width of the cathodic pulse, we can determine the charge injection capacity (Q_inj_).

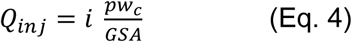

### Optical Transparency – UV-Vis-IR Spectroscopy

To characterize the optical transparency of Smart Dura, we performed UV-Vis-IR spectroscopy analysis using the setup shown in Fig. 4A. Using an incandescent light bulb to generate a broadband light source, we illuminated a large area of the Smart Dura from the bottom via a 45-degree broadband aluminum mirror (Thorlabs, CCM1-F01) and a ring-shaped holder (Thorlabs, CP1XY). An optical fiber (200 µm core diameter glass fiber, QP200-2-SR-BI) coupled to an optical spectrometer (Ocean Optics HR4000CG-UV-VIS) was used to measure the transmission spectrum. The fiber was placed directly above the Smart Dura surface and inside the PDMS chamber walls. The disk-shaped PDMS chamber was used to hold the water for the Smart Dura and water transmission measurement. We first measured the optical transmission of Smart Dura through air. Then, water was carefully added with a syringe inside the PDMS chamber to make sure the fiber tip was immersed in water to characterize the optical transmission through the Smart Dura and water. Prior to placing the Smart Dura on top of the holder, we performed measurements with the Smart Dura removed from the light path to obtain measurements with the light source turned off (dark reading) as well as turned on (air reference reading). The dark reading is used to offset the effects of any coupled light into the fiber from the room environment. The reference reading is used to obtain a baseline measurement with no sample. With all three measurements: dark, reference, and sample readings, we determine the optical transmission of the Smart Dura by first subtracting the dark reading from both reference and sample readings and then taking the ratio between the sample and reference measurements. This was repeated for the Smart Dura and water transmission measurement.

### Solid Mechanics Finite Element Analysis (FEA)

To study the mechanical performance of our Smart Dura, we performed from finite element analysis (FEA) solid mechanics simulations using COMSOL Multiphysics. In this simulation study we simulated a membrane with the same geometry as the Smart Dura (20 mm in diameter and 260 μm total thickness) made out of two different PDMS + Parylene C bi-layer configurations:

i. Bilayer PDMS (250 μm thick) + Parylene C (10 μm thick)
ii. Bilayer PDMS (220 μm thick) + Parylene C (40 μm thick).

For both configurations, the membrane is assumed to be fixed on its edges, with an applied uniform load pressure applied from the bottom of the membrane with values typical of CSF pressures in primates (1-10 mmHg or 133.32-1333.2 Pa). We used a linear elastic mechanical model for Parylene C and a Neo-Hookean hyperelastic mechanical model for PDMS. All material properties were extracted from the material library provided by COMSOL.

### Animals and Data Acquisition

Animal care and experiments were approved by the University of Washington’s Office of Animal Welfare, the Institutional Animal Care and Use Committee, and the Washington National Primate Research Center (WaNPRC). We validated our Smart Dura on three male macaques

(monkey M: *Macaca nemestrina*, 18.04 kg, 20 years; monkey N: *Macaca nemestrina*, 4.75 kg, 4 years, monkey H: *Macaca mulatta*, 13.2 kg, 11 years). Monkeys M and N underwent craniotomy and durotomy targeting the sensorimotor and visual cortices, respectively. Monkey H has a pre-established chronic cranial window overlying the left posterior parietal cortex (PPC)^52,53^. The details of our surgical procedure are described here^45^.

All electrophysiological recordings were performed with Grapevine Nomad processors, Front Ends (FEs), and Trellis software (Ripple Neuro, UT) in combination with our Smart Dura system. Two flexible PCBs of individual Smart Dura were connected to rigid PCBs and secured on our customized holder. Each rigid PCB was then attached to the FE (Nano2+Stim), interfacing with the Grapevine Nomad processor, to filter, amplify, and digitize the neural signals. Raw signals were recorded at a sampling rate of 30 kHz with a bandpass filter between 1 Hz and 7.5 kHz using the Trellis software. LFPs were sampled at 1 kHz by digitally filtering using a low-pass filter of 250 Hz with a notch filter at 60, 120, and 180 to remove line noise. Spikes were detected by high-pass filtering with a cutoff frequency of 250 Hz and setting the threshold to minus six standard deviations of the background noise. Spike sorting of the detected spikes was performed by K-means clustering using Offline Sorter software (Plexon Inc., TX). Subsequent signal processing and data analyses were conducted using MATLAB (MathWorks, MA).

The empirical SNR of the spikes detected and shown in figure 6 was calculated with the following equations:

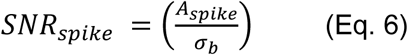

where 𝐴_𝑠𝑝𝑖𝑘𝑒_ is the amplitude of the mean spike waveform and 𝜎_𝑏_ is the standard deviation of the background noise. For an empirical SNR value in decibels, we use equation 7 below:

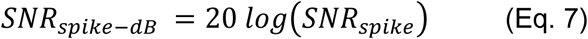

### Optical Transparency – Two-photon imaging

To validate the optical transparency of the Smart Dura, we conducted *in vivo* vascular imaging in monkey N under anesthesia using two-photon microscopy. A 64-channel Smart Dura was placed on the left hemisphere of the visual cortex, and FITC-dextran (50 mg/kg mixed at 50 mg/mL in sterile water for injection; Sigma-Aldrich) was injected intravenously. Two-photon imaging was performed with a multi-photon microscope (MOM; Sutter Instrument) equipped with a Ti:sapphire laser set to 920 nm wavelength (Chameleon Vision II; Coherent). The excitation laser beam was focused with a 16x, NA 0.8 water immersion objective lens (CFI75 LWD 16X W; Nikon). The microscope was controlled by ScanImage software (Vidrio Technologies).

### Behavior experiments and Data analysis

To characterize the Smart Dura’s ability to record movement-related neural signals, we custom-designed and trained monkey H to perform a MATLAB-based, center-out-reaching task similar to the one discussed in our previous work^54^. During these behavioral sessions, the animal was head-fixed and seated in a primate chair in front of the monitor inside our experimental rig. For each session, a reflective sticker was placed on the middle finger of the monkey’s right hand, contralateral to the implanted cortex. The sticker was tracked by a motion capture system (Motive, OptiTrack; NaturalPoint Inc., OR) to estimate finger location and movement trajectory. The tracking data was streamed and processed through MATLAB in real time to control the task directly. During each trial, the monkey was instructed to hold in the center green circle until the go-tone and then move its finger to the red target that would appear at one of four possible locations on the screen. For each successful trial, the animal was rewarded with juice.

We recorded neural activity using 32-channel and 64-channel arrays for epidural recordings and a 64-channel array for subdural recordings. LFPs were preprocessed to remove noise and artifacts. This included band-pass filtering to isolate the relevant frequency bands and manual inspection to exclude channels with excessive noise and segments with large artifacts. High-impedance electrodes were also excluded from subsequent analysis. For LFP power analyses, we computed signal power by squaring the signal and dividing the sum by its duration and focused on the theta (4-8 Hz) and gamma (30-70 Hz) frequency bands, which are known to be involved in motor control and cognitive processes.

For the epidural recordings, we analyzed the power changes in the theta band during both task and resting periods. During the task period, we first set the segments of interest to span the entire trial, from the trial start (start target being shown) to the successful completion of the trial (animal holding the end target). This task period is usually about 2000 ms in length, depending on the specific trial. LFPs recorded during the no-movement period before each trial (after the previous trial was completed) were used as a baseline to calculate task-related power changes. Next, we performed a similar power analysis on resting period recordings to serve as controls. Resting baseline recordings either before or after the task sessions were randomly selected to match the length of individual trials during the reach task. For both task and resting data, we visualized trial-averaged power changes from baseline using heatmaps for each good electrode and interpolation between electrodes and plotted histograms for selected individual channels to show the distribution of signal power across trials for both task and resting periods. For subdural recordings, we extended the analysis to both the theta and gamma bands during the task and resting periods, using the same procedure as for the epidural recordings, allowing us to compare power changes across different frequency bands.

To examine the time-frequency distribution of the subdural LFPs during the reach task, we generated multitaper spectrograms that were z-scored to the 500 ms trial segment before the go-tone, providing a trial-averaged time-frequency spectrum for the period from 500 ms before to 1000 ms after the go-tone. Additionally, to compare the LFP powers at individual frequency bands, we computed the average power of the subdural LFPs before and after the go-tone in the theta (4-8 Hz), alpha (8-15 Hz), beta (15-30 Hz), and gamma (30-70 Hz) bands and assessed statistical significance using paired t-tests with post hoc Bonferroni corrections for multiple comparisons (family-wise error rate of 0.05).

### Tactile Stimulation and Data Analysis

We performed tactile stimulation in monkey M during anesthesia under urethane (0.52 g/L). Using a custom-made tactile vibrating stimulator and a code implemented in MATLAB, 30 stimulation pulses of 1 second were delivered to the monkey’s left-hand fingertips (Fig. 8A, B). For neural activity recording, a 64-channel Smart Dura was placed on the right hemisphere of the sensorimotor cortex. LFPs were low-pass filtered with a 40 Hz cutoff frequency. To assess evoked responses during the tactile stimulation, we calculated power 100 ms before and 200 ms after the stimulus onset for each trial by taking the sum of the squared signal divided by its duration. For quantitative comparison, a two-sample t-test was performed. Spectrograms for individual electrodes were obtained by generating a multitaper spectrogram (30 ms time window, 3 ms overlap) for each trial and then averaging over all trials. The spectrograms were z-scored with respect to the neural activity 500 ms preceding the stimulus, and the power in the 1-40 Hz frequency band was averaged at each time point.

### Epidural Electrical Stimulation and Data Analysis

To validate the electrical stimulation capability of the Smart Dura, we applied electrical stimulation in a paired stimulation paradigm with epidural neural recording in monkey H. Throughout each recording session, the animal was head-fixed in a primate chair watching an animated children’s television show while receiving variable amounts of juice at random intervals, and a 64-channel Smart Dura was set on top of the dura mater over the left PPC. Each block for recording and stimulation consisted of two consecutive 10-minute spontaneous LFP recordings (‘Baseline 1’ and ‘Baseline 2’ in Fig. 9A), followed by a 10-minute electrical stimulation on two unique cortical sites (‘Stimulation’ in Fig. 9A), and a 10-minute post-stimulation LFP recording (‘Post-Stimulation’ in Fig. 9A). During each stimulation period, 5 Hz burst stimuli were delivered through two randomly chosen electrodes with a 10 ms delay. A burst for each site was comprised of 5 charge-balanced biphasic pulses at a pulse frequency of 1 kHz with a pulse width of 200 µs and an amplitude of 5 µA. A total of 12 blocks were recorded.

To monitor changes in local activity, we obtained low gamma (30-59 Hz) and high gamma (60-150 Hz) band signals via bandpass filtering. Artifacts in the signal were excluded from further analysis by normalizing the signal and identifying samples in which the absolute value of the normalized signal exceeded 10 standard deviations from the mean. After removing artifacts, the signal recorded from each channel was normalized to the mean and standard deviation of the first baseline period. Next, we calculated the signal power of each 10-minute period by squaring the signal and dividing the sum by the time duration in seconds. To assess differences in spontaneous versus stimulation-induced changes in power, we compared the power changes between the two initial baseline periods and between the post-stimulation and the first baseline periods and performed paired sample t-tests with post hoc Bonferroni corrections for multiple comparisons (a family-wise error rate of 0.05).

### Optogenetic stimulation

For viral vector delivery and opsin expression, we performed convection-enhanced delivery ^28^ of a red-shifted inhibitory opsin Jaws (rAAV8-hSyn-Jaws-KGC-GFP-ER2; UNC Vector Core, NC) into the PPC of monkey H during the surgery for a chronic cranial window^52,53^ . To activate the opsin, we used a 638 nm laser (Doric Lenses, Quebec, Canada) coupled with an optical fiber and placed the optical fiber over the Smart Dura. 30 light pulses of a laser power (a beam radius of about 0.7 mm ^28^) were delivered for 900 ms every 5 seconds. To obtain a heatmap, multitaper spectrograms (150-ms time window with 3-ms steps) were computed and normalized by z-scoring with respect to the frequency band power 350 ms before the onset of stimulation. The bins between 4-40 Hz during the stimulation were averaged and divided by electrode impedance to eliminate the influence of impedance differences between electrodes. The evoked response was considered significant if the averaged power during stimulation was significantly greater (§p < 0.05) using a one-sample t-test, where the null hypothesis is rejected when the mean is not equal to 0.

## Acknowledgements

This work was supported by the National Institute of Neurological Disorders and Stroke of the National Institutes of Health under award number R01NS116464, R01NS119395, and U01NS115585, the National Institute of Biomedical Imaging and Bioengineering of the National Institutes of Health under award number EB029365, the National Institute of Mental Health of the National Institutes of Health under award number R01MH125429 and UG3MH126864, the National Institutes of Health T32 training in theoretical and computational approaches to neural circuits of cognition under award number 1T32MH132518 and the National Institutes of Health Blueprint and BRAIN Initiative D-SPAN Award 1F99NS130828. The content is solely the responsibility of the authors and does not necessarily represent the official views of the National Institutes of Health. This work was also supported by the Washington National Primate Research Center P51 OD010425 and U42 OD011123 from the National Institutes of Health Office of Research Infrastructure Programs, the Washington Research Foundation, the American Heart Association, the Weill Neurohub, and the National Science Foundation Graduate Research Fellowship Program. The authors acknowledge the use of the Bertucci Nanotechnology Laboratory at Carnegie Mellon University supported by grant BNL-78657879. We thank the staff of the Washington National Primate Research Center for their help with animal care.

---

**Figure S1.**
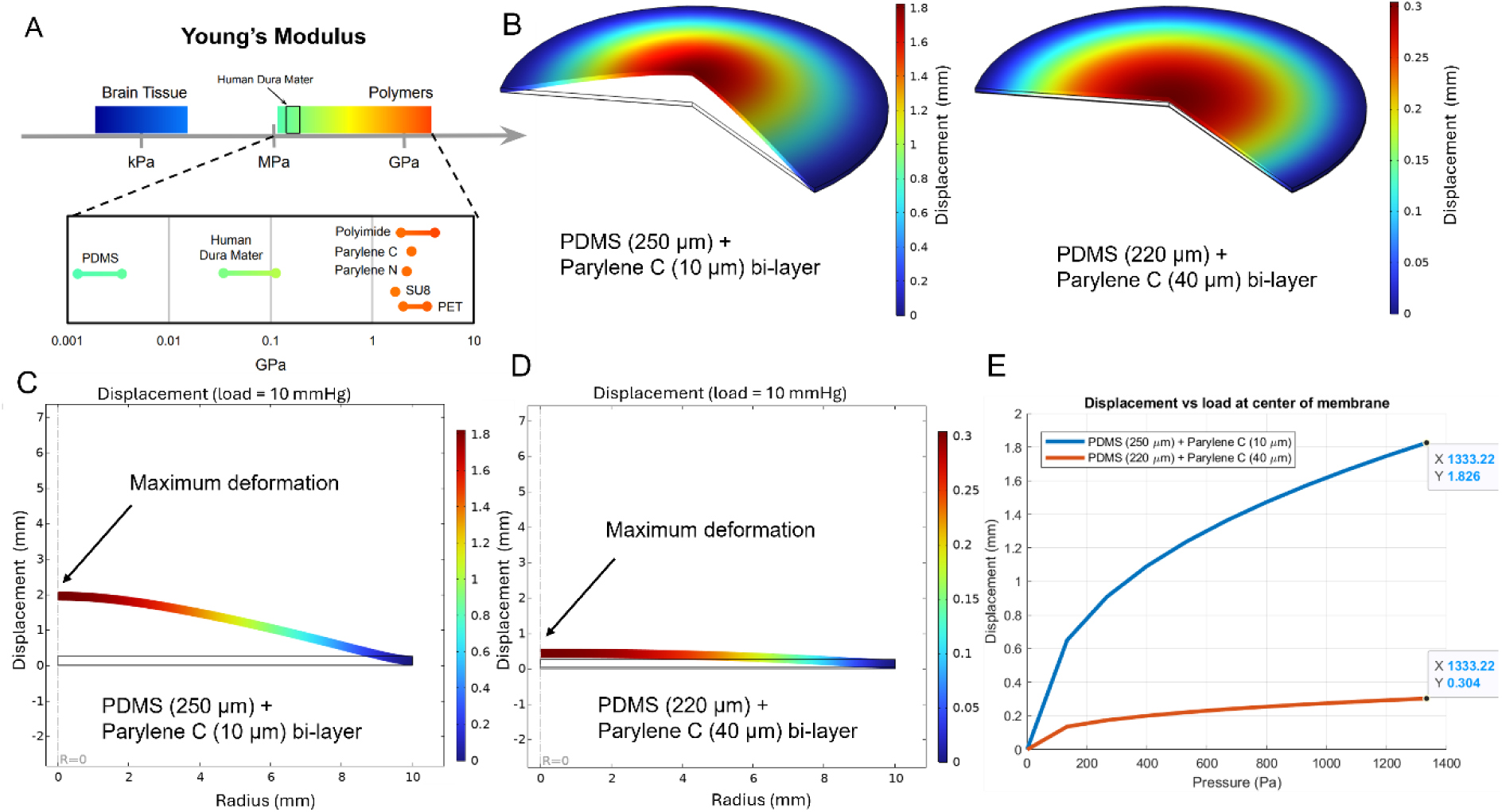
Material choice and tunable mechanical properties of Smart Dura. A) Young’s moduli of polymers used for neural interfaces, brain tissue, and native dura. Figure modified with permission from [^25^]. B) Comparison of the 3D deformation profile of 250 µm and 10 µm (left) and 220 µm and 40 µm (right) PDMS/Parylene C bi-layer substrates with 10 mmHg uniform pressure load applied to the bottom. C) Cross-section 2D plot of the deformation from the center of the membrane to its edge for the 250 µm and 10 µm PDMS-Parylene C bi-layer. D) Cross-section 2D plot of the deformation from the center of the membrane to its edge for the 220 µm and 40 µm PDMS-Parylene C bi-layer. Note the notable difference in deformation compared to C). E) Comparison of displacement at the center of the membrane of the two difference thickness ratio PDMS-Parylene C bi-layers subject to different nominal cranial pressure loads (1-10 mmHg or 133.32-1333.2 Pa), showing the capability of tuning the mechanical stiffness of the bulk bi-layer substrate based on the relative thickness ratio of PDMS and Parylene C.

**Figure S2.**
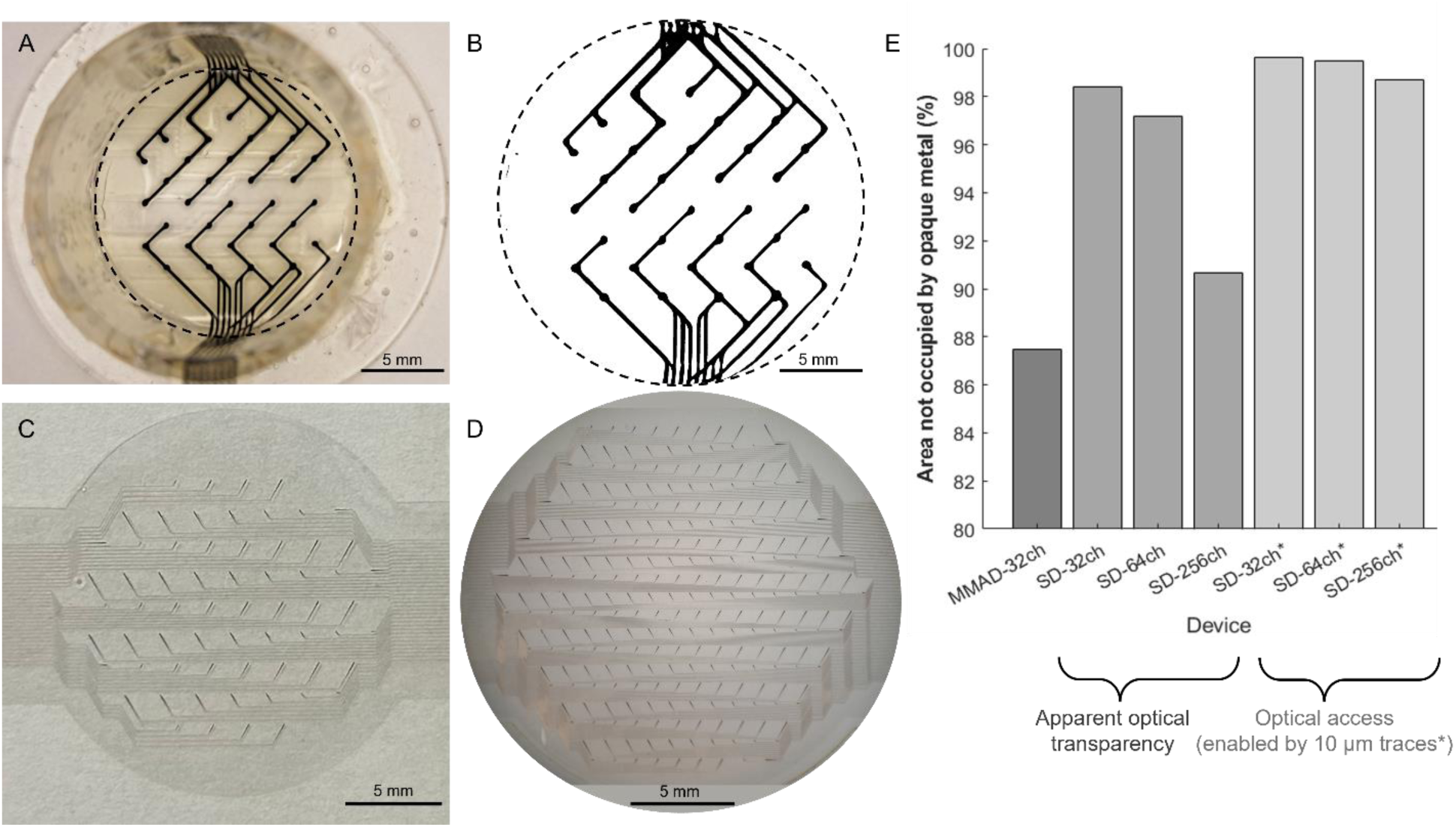
Comparison of optical window areas. A) Photograph of 32-channel MMAD with 0.5 mm diameter electrodes and 250-300 μm wide traces. B) Binary image of the optical window of the MMAD used to calculate the percentage of white (transparent) vs black (opaque) pixels to obtain the percentage of area of the MMAD optical window not blocked by the opaque metals and electrodes. C) Photograph of 64-channel Smart Dura over a white background. D) Photograph of the high-density 256-channel Smart Dura over a white background. E) Bar plot comparing the percentage of the optical window area not blocked by opaque electrodes and traces, where SD stands for Smart Dura. The 32-channel MMAD has an optical access of 87.46% while the Smart Dura has optical access for the 32-channel, 64-channel, and 256-channel of 98.42%, 97.19%, and 90.68% respectively, demonstrating the improved optical access of the Smart Dura provided by the microfabrication techniques described in this paper. If 10 μm trace width features are further neglected in Smart Dura designs as discussed above, the optical access for the 32-channel, 64-channel, and 256-channel increases to 99.64%, 99.46%, and 98.69% respectively.

**Figure S3.**
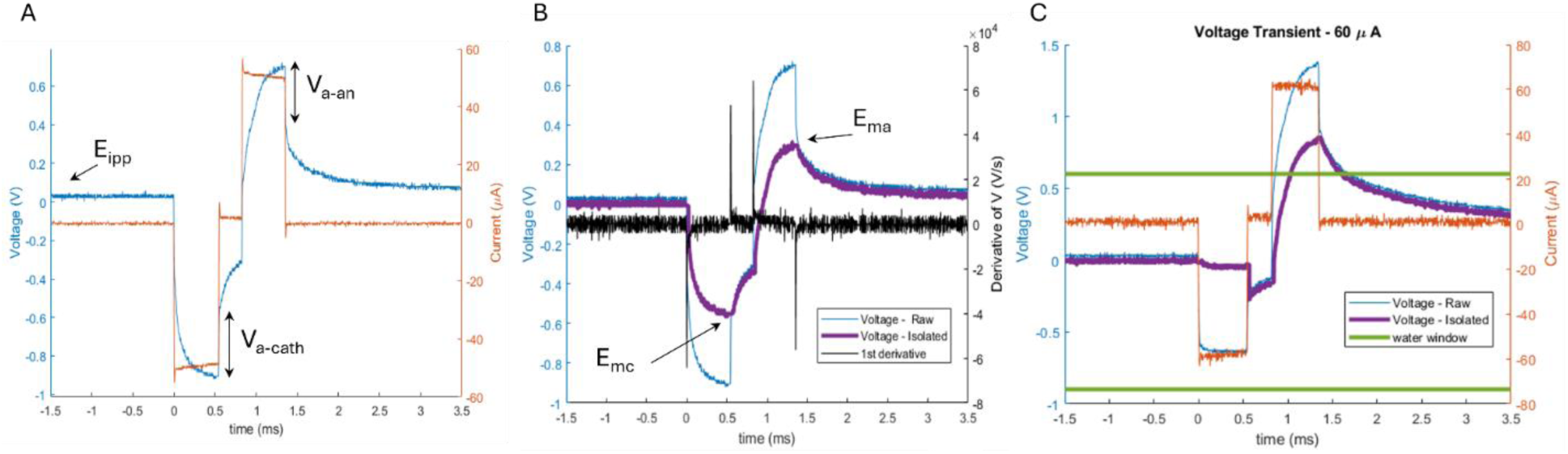
Illustration of the process to obtain the isolated voltage transient waveform across the electrode-electrolyte interface from raw measurements to accurately calculate the charge injection capacity of the micro-electrodes from Smart Dura devices. A) Raw data from biphasic current pulse (red) and resulting voltage transient waveform recorded at the electrolyte from the Ag-AgCl reference electrode. B) First derivative (black) of raw voltage transient waveform (blue) used to identify the location of pulse edges (t_Va_) from peaks to isolate the voltage exclusively across the electrode-electrolyte (cyan).

